# Hormonal control of motivational circuitry orchestrates the transition to sexuality in Drosophila

**DOI:** 10.1101/852335

**Authors:** Stephen X. Zhang, Ethan H. Glantz, Dragana Rogulja, Michael A. Crickmore

## Abstract

Newborns and hatchlings of many species perform incredibly sophisticated behaviors, but all vertebrates and many invertebrates selectively abstain from sexual activity at the beginning of life. Hormonal changes have long been associated with adolescence, but it is not clear how these circulating factors create a new motivation and drive its associated behaviors. We show that the transition to sexuality in male *Drosophila* is controlled by juvenile hormone, which spikes at eclosion and declines over days as the propensity for courtship gradually increases. Juvenile hormone directly inhibits the activity of at least three courtship-motivating circuit elements, ensuring the complete suppression of sexual motivation and behavior. Blocking or overriding these inhibitory mechanisms evokes immediate and robust sexual behavior from very young and otherwise asexual males. These results provide a first example of hormonal changes gating the transition to sexuality by activating latent, but largely developed and functional, motivational circuitry.

## INTRODUCTION

The intensities of all motivations vary with time and circumstance, but the complete absence of mating behaviors in young animals suggests the non-existence of reproductive drive until later in life. In mammals, the emergence of sexual behavior is associated with changes in circulating hormones, but it is not clear whether these hormones awaken a latent drive or instruct the brain to build the foundation for a new one. This question has been most commonly considered under the organizational-activational hypothesis (Phoenix et al., 1959). Organizational functions are long lasting and structural, such as the prenatal establishment of sexual identity and the maturation of brain circuitry during adolescence (e.g., Simerly, 1989), while activational functions use previously organized neuronal circuitry to promote sexually dimorphic behaviors and tendencies (Wu et al., 2009). Despite this longstanding framework, the extent to which the motivational component of the transition to sexuality results from either organizational or activational hormonal functions is not clear (Schulz et al., 2009).

Indirect evidence for an organizational role of gonadal hormones in the onset of sexual behavior comes from the long timescales over which their effects become apparent (Sisk and Zehr, 2005). For example, administration of testosterone to juvenile or castrated adult male rodents induces sexual behavior, but only after several days or weeks (Putnam et al., 2001; Södersten et al., 1977), in accord with the delay between testosterone elevation and the onset of sexuality at puberty (Sisk and Zehr, 2005). Similarly, while castration immediately causes a large reduction in circulating testosterone, the resulting progression towards asexuality is gradual (Davidson, 1966; Krey and Mcginnis, 1990). To our knowledge, no experimental evocation of juvenile sexual behavior has been reported in any animal without manipulation of hormonal signaling, which might be taken as further evidence for organizational gating of sex drive. However, the circuitry underlying sexual motivation and behavior is poorly understood in almost all animals, preventing a thorough analysis of its potential functionality during asexual stages of life.

Recently we and others have characterized much of the circuitry underlying the decision of male *Drosophila* to court females (Clowney et al., 2015; Kallman et al., 2015) and how this decision is weighted by the male’s recent mating history (Zhang et al., 2016, 2018, 2019). Since newly eclosed male flies never engage in courtship (Spieth, 1974), this system presents a unique opportunity to examine the molecular- and circuit-level mechanisms underlying the emergence of sexual motivation. Though in the fly each cell’s sexual identity is independently determined by cell-intrinsic mechanisms (Baker and Belote, 1983), organism-wide developmental transitions (e.g. metamorphosis) are organized by hormonal signaling (Mirth et al., 2014). Here we show that hormonal changes also control the transition to sexuality, and that sexual behavior emerges from the relaxation of stringent inhibition imposed on developed, but functionally dormant, motivational circuitry.

The identification and evaluation of a potential mating partner requires multisensory integration, but in our experimental paradigm the decision to court is ultimately triggered by a male tapping a female with his pheromone-receptor-bearing leg (Fan et al., 2013; Kohatsu and Yamamoto, 2015; Kohatsu et al., 2011; Thistle et al., 2012). The tap delivers parallel excitatory and inhibitory inputs to the male’s P1 courtship command neurons which initiate and maintain courtship when sufficiently stimulated (Clowney et al., 2015; Kallman et al., 2015; Zhang et al., 2018). The sensitivity of P1 neurons to the inhibitory input from a tap is decreased by a local dopamine signal, which, in mature males, is tuned to reflect recent mating history (Zhang et al., 2016). If a male has not mated for several days, the dopamine tone is high, increasing the probability that a tap will lead to courtship. Once courtship has commenced, the same dopamine signal maintains courtship bouts for the tens of seconds to several minutes required before mating commences. This motivating dopaminergic activity is maintained for days in the absence of females, is decremented by each mating, and slowly recovers over 3-4 days.

Matings promote satiety by activating a set of Copulation Reporting Neurons (CRNs). These neurons project dendrites to the external genitalia and send axons to the brain, where they reduce the activity of Fruitless-positive NPF neurons. This decrease is relayed through the NPF Receptor to the motivation-promoting dopamine neurons, reducing their activity. After a few matings, substantial satiety is induced, and reproductive motivation remains low for several days. Mating drive has an intrinsic tendency to recover due to recurrent excitation: NPF neurons excite, and are excited by, Doublesex-positive pCd neurons (Zhang et al., 2019), forming a loop that holds and gradually accumulates activity during periods of abstinence. But loop activity is prevented from immediately rebounding by the prior activity of the transcription factor CREB2, which, during the period of high motivation that precedes mating, transcribes inhibitory genes (e.g. potassium channels) that will sustain the decremented activity state, forcing recovery to proceed on a biochemical, rather than electrical, timescale (Zhang et al., 2019).

This circuit architecture (see Figure 3A) suggests many hypotheses for the prevention of courtship in newly eclosed males. For example, a juvenile male might not recognize or tap females; or the CRNs may be constantly active, inducing overwhelming satiety; or, as in the organizational hypothesis, some or all of the courtship circuitry may not be fully developed. We present evidence refuting all these hypotheses. Instead, we find that a spike in juvenile hormone levels at eclosion directly and selectively imposes long-lasting activity suppression on all known motivation-promoting circuit elements: both populations of loop neurons as well as the downstream dopaminergic neurons. Overriding any of these repressive mechanisms evokes robust courtship in extremely young males, providing clear evidence that the underlying circuitry is functional, but dormant, in juveniles. Multi-tiered suppression of motivational circuitry appears to be the key difference between the complete inactivation of mating drive in early life and its modulation thereafter.

## RESULTS

### A latent capacity to court in juvenile males

Male flies show no courtship behavior during the first 6 hours after eclosion, a stage we refer to as juvenile (Spieth, 1974) (Figures 1A and 1B). Sexual behavior then gradually increases, such that ∼90% of males will show some courtship toward virgin females by 24 hours after eclosion (Figure S1A). Early courtship bouts terminate frequently, but they become longer with maturation such that males achieve a near peak courtship index (fraction of time spent courting over 5 minutes) by 72 hours (Figures 1B and S1A). This increasing courtship behavior mirrors gains in reproductive potency, as measured by the reserves of sperm and seminal fluid stored in the ejaculatory bulb (Figures 1A and 1C) and resembles the gradual and initially fragmented recovery from satiety seen in mature males (Figures S1B-S1C) (Zhang et al., 2018). In mature males, the reproductive organs themselves have no apparent impact on mating drive (Zhang et al., 2016), and, as expected, their removal from juveniles did not disinhibit courtship (Figure S1D).

**Figure 1.**
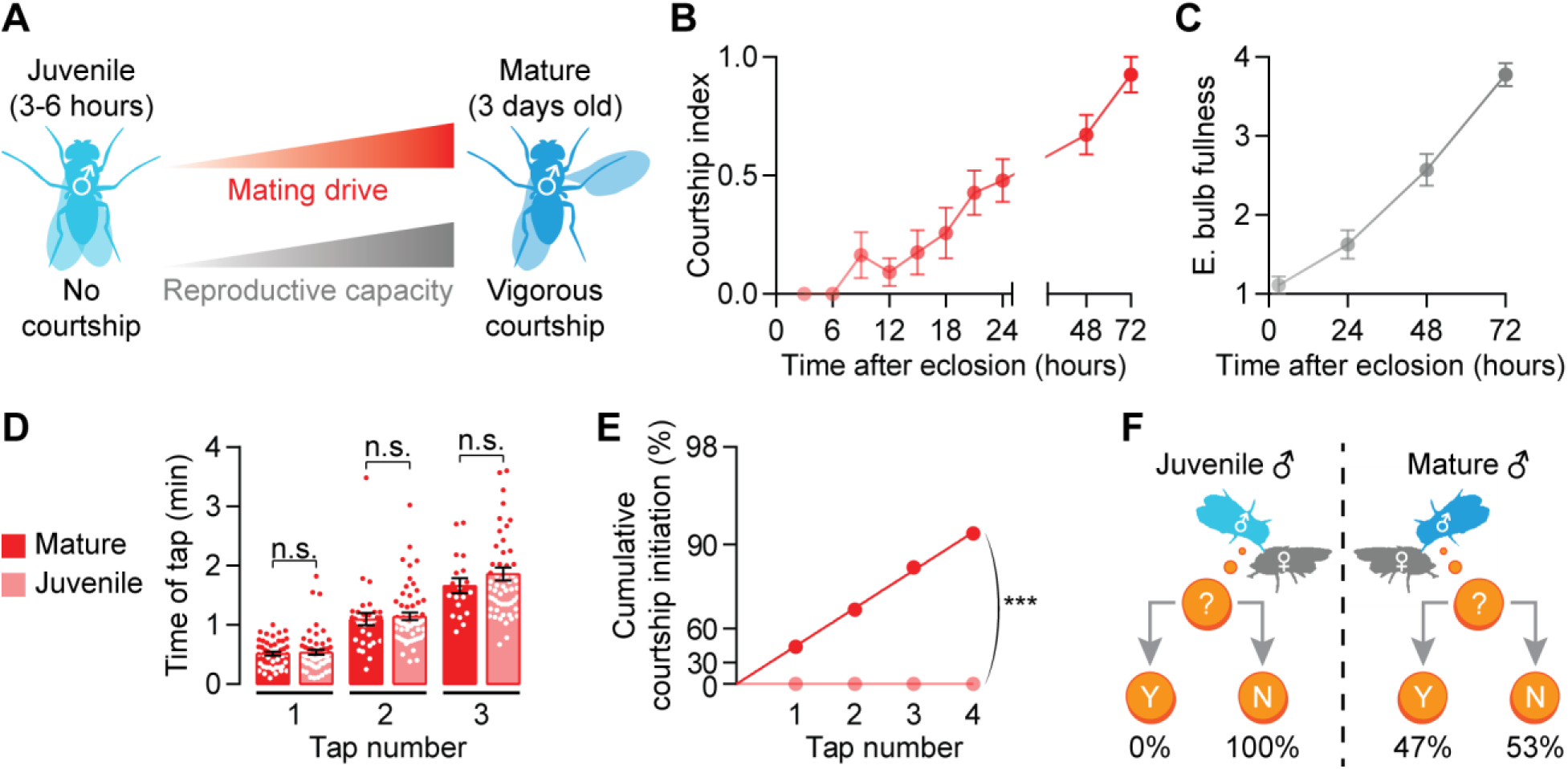
Male flies gradually accumulate mating drive and reproductive capacity after eclosion. (A) Schematic showing accumulating mating drive and reproductive capacity from juvenile (3-6 hours after eclosion) to mature (3 days) male flies. (B-C) Both courtship and ejaculatory bulb fullness increase steadily over the first 3 days after eclosion (B: n = 12-32 males, C: n = 7-9^†^). (D) Mature and juvenile males tap females with similar frequency (one-way ANOVA^‡^, n = 48-57 males). (E-F) Each tap by mature males has a ∼47% chance of initiating courtship, whereas taps by juveniles never trigger courtship (bootstrap, n = 48-57 males). ^†^Throughout the paper, error bars represent SEM unless otherwise stated, but are left out of tap-induced courtship plots because of log-scaling. ^‡^***p<0.001, **p<0.01, *<0.05, n.s. not significant for all figures.

In our experiments (e.g. Zhang et al., 2018), each courtship bout is initiated by the male tapping the female with a pheromone-receptor-bearing leg, and, in mature males, each tap triggers courtship with a probability that reflects reproductive capacity. We found no difference in the frequency with which juvenile males tapped mature females (Figure 1D), but juvenile taps never led to courtship (Figures 1E-1F). The suppression of sexual behavior in juveniles is therefore independent of the decision to collect sensory information by tapping and instead must occur somewhere in the circuitry that translates this information into courtship behavior.

A set of ∼20 P1 neurons per brain hemisphere (Figure 2A) is the primary decision center for courtship initiation and maintenance: their inhibition or feminization prevents courtship (Kimura et al., 2008; von Philipsborn et al., 2011; Zhang et al., 2018), and their stimulation evokes courtship regardless of the male’s satiety state or the quality of the courtship target (Pan et al., 2012; von Philipsborn et al., 2011; Zhang et al., 2016). We found no obvious anatomical differences between the P1 neurons of mature and juvenile males (Figure 2A). Strikingly, thermogenetic stimulation of juvenile P1 neurons using the warmth-sensitive cation channel TrpA1 evoked robust courtship, including orienting toward the female, following, and singing to her (Figure 2B). This courtship did not escalate to abdomen bending for copulation initiation, potentially due to incomplete development of abdominal muscles (Lawrence and Johnston, 1986) or hardening of the abdominal cuticle. These results show that the neural circuitry responsible for executing most courtship behaviors is developed and capable of functioning in juvenile males, implying that the prevention of courtship must occur somewhere downstream of a tap and upstream of P1 activation.

**Figure 2.**
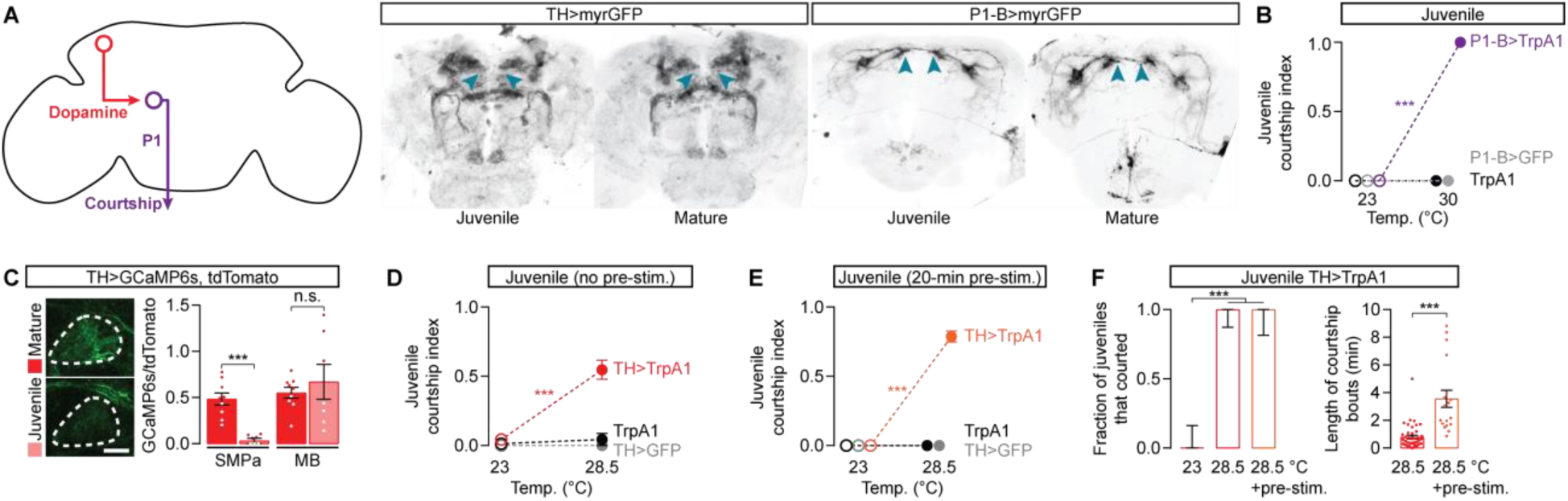
Mating-drive circuitry is functional but quiescent in juveniles. (A) Left: schematic depicting dopamine-to-P1 circuitry. Right: dopaminergic and P1 neurons are morphologically similar in juvenile and mature males. Blue arrowheads point to the SMPa. (B) Thermogenetic stimulation of courtship commanding P1 neurons evokes courtship from juvenile males (two-way ANOVA, n = 5-9 males). (C) In juvenile males, calcium activity is low in dopaminergic projections to the SMPa but not in the neighboring vertical lobe of the mushroom body (one-way ANOVA, n = 7-9 male brains). (D-E) Thermogenetic activation of juvenile dopaminergic neurons drives courtship (D), which becomes more vigorous in males that have been pre-stimulated for 20 minutes prior to the assay (two-way ANOVA, D: n = 8-31 males, E: n = 12-15). (F) Acute dopaminergic stimulation causes frequent courtship initiation (left), but consolidated courtship bouts require prolonged stimulation (right) (Left: Fisher’s exact test, n = 12-17 males; Right: t-test, n = 14-43 bouts from 12-17 males).

**Figure 3.**
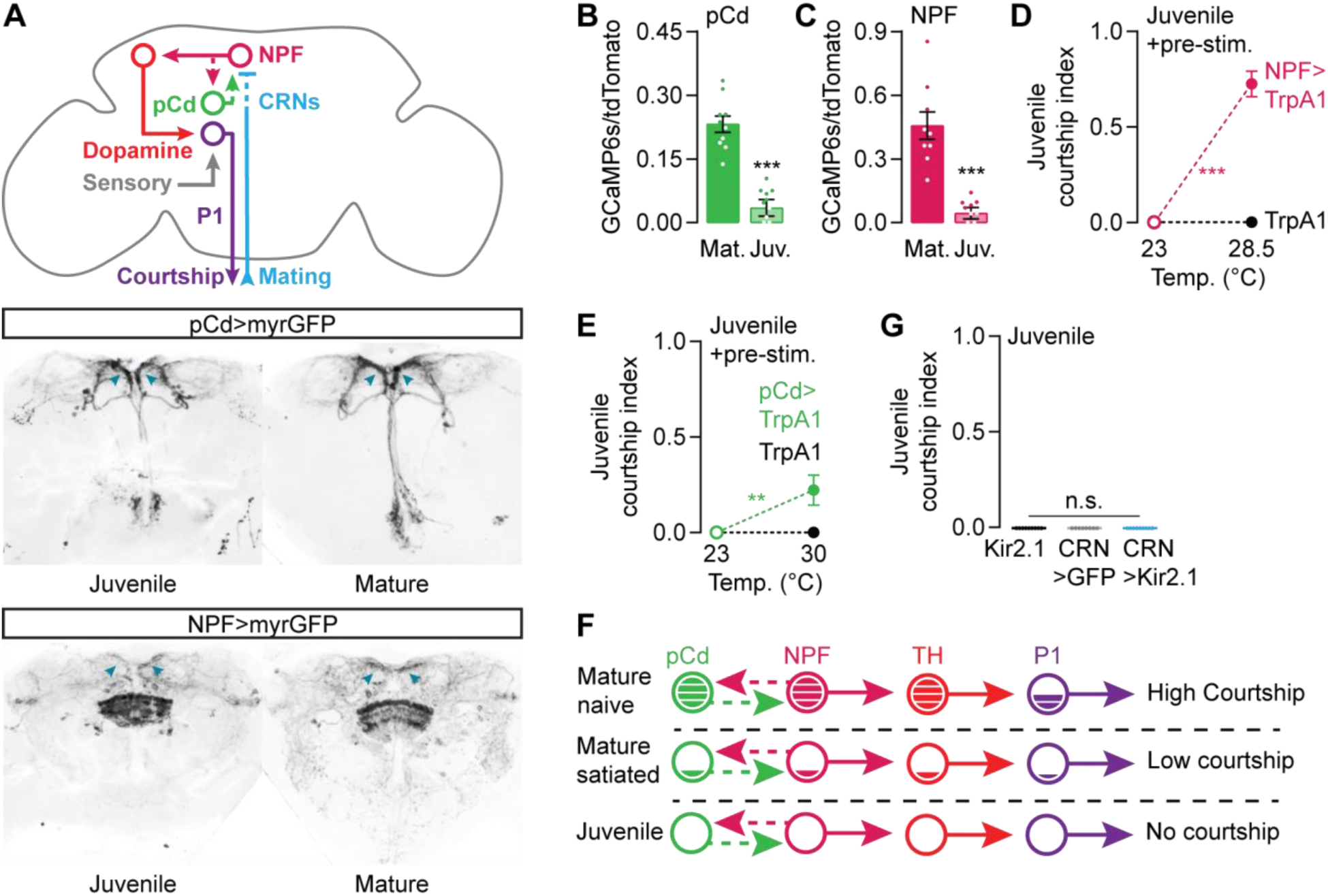
Upstream inputs to the motivating dopamine neurons are quiescent in juvenile males. (A) Left: schematic depicting mating-drive circuitry, solid lines between circuit elements indicate connections that are known to be direct. Right: expression patterns of 41A01-Gal4 (pCd) and NPF-Gal4. (B-C) Baseline calcium activities in the SMPa for both pCd (B) and NPF neurons (C) are low in juvenile males (t-test, B: n = 10 brains each, C: n = 9 each). (D-E) Thermogenetic stimulation of NPF (D) or pCd (E) neurons beginning 20 minutes before a courtship assay induces courtship from juvenile males (two-way ANOVA, D: n = 14-16 males, E: n = 13-17 males, see Methods for explanation of stimulation temperatures). (F) Mating-drive circuitry is quiescent in juveniles. (G) Silencing the satiety-inducing CRNs does not promote courtship in juveniles (one-way ANOVA, n = 8-9 males).

### Stimulating motivational circuitry evokes courtship in juveniles

Dopaminergic activity in the anterior of the superior medial protocerebrum (SMPa) is a functional neural correlate of mating drive: it is high in motivated males and decreases with satiety to reduce courtship behavior (Zhang et al., 2016). This dopaminergic signal is received by P1 where it sets the probability of courtship initiation following a tap (Zhang et al., 2018). In juveniles, dopaminergic projections to the SMPa are evident (Figure 2A), but the baseline activity of these projections is essentially undetectable, though nearby mushroom-body-projecting dopaminergic neurons showed high levels of activity (Figure 2C). When we thermogenetically stimulated either all dopaminergic neurons (Figure 2D), or only the courtship-motivating subset (Figures S2A-S2B) (Zhang et al., 2016), we induced robust courtship from juvenile males. This courtship was qualitatively similar to that evoked by P1 stimulation, though the males spent only ∼50% of their time courting, compared to ∼100% with P1 stimulation. In previous work on this circuitry (Zhang et al., 2018), we found that while acute dopaminergic stimulation evokes frequent courtship initiation from satiated mature males, P1 must receive prolonged dopaminergic signaling before courtship bouts become sustained. Similarly, stimulation of juvenile dopaminergic neurons that began 20 minutes before presentation of a female resulted in longer courtship bouts and a near maximal courtship index (Figures 2E-2F and S2A-S2B). These results show that, similar to satiated mature males, the lack of dopaminergic activity in the SMPa is causal for the asexual behavior of juveniles. As dopaminergic activity increases with age, or through synthetic stimulation, males first increase their propensity to initiate courtship and then gradually consolidate these bouts for the length of time often required to achieve copulation (Figures S1A and S1C).

The motivating dopamine tone in mature males is set by the activity held and adjusted within a recurrent excitation loop. The loop contains two known populations of neurons: Fruitless-positive NPF neurons that directly stimulate the dopaminergic neurons, and Doublesex-positive pCd neurons that are stimulated by, and feedback to stimulate, the NPF neurons (Figure 3A) (Zhang et al., 2019). In juveniles, the morphologies of NPF and pCd neurons appear normal, including their projections to the SMPa (Figure 3A). As with the dopaminergic neurons, the activities of both NPF and pCd populations are high in mature, sexually naïve males (Zhang et al., 2019), but extremely low in juvenile males (Figures 3B and 3C). Stimulation of the NPF neurons drives juvenile courtship to a similar extent as direct activation of the dopaminergic neurons (Figure 3D and S2C). Stimulation of the pCd neurons also induced juvenile courtship (Figure 3E), though less effectively (Figure S2D), presumably due to their indirect access to the dopaminergic neurons (Zhang et al., 2019). In summary, these results reveal a qualitative similarity between the motivational circuitries of highly satiated mature males and juvenile males (Figure 3F), though the quantitatively stronger suppression in juveniles implies additional mechanisms.

During mating, satiety is incrementally induced in the male through the activity of the Copulation Reporting Neurons (CRNs) (Figure 3A), which project dendrites to the bristles surrounding the external genitalia and report copulations by recruiting inhibition onto NPF neurons in the brain (Zhang et al., 2019). In mature males, silencing the CRNs during mating prevents satiety, while stimulating them in males that have never mated induces lasting satiety (Zhang et al., 2019). Like the motivational circuitry examined above, the CRNs were capable of functioning in early life: CRN stimulation produced a long-lasting delay in the gradual increase of courtship behavior (Figures S2E and S2F). However, CRN activity is dispensable for the suppression of juvenile sexual behavior, as their tonic silencing using the potassium leak channel Kir2.1 did not provoke courtship from juveniles (Figure 3G). This result rules out the hypothesis that the CRNs suppress sexual behavior in juveniles, pointing to the existence of an alternative mechanism for preventing the accrual of mating drive in young males.

### A decreasing juvenile hormone titer gradually releases mating drive

In mature males, the enduring effects of prior motivation sustain satiety after mating (Zhang et al., 2019). How might the intrinsic tendency for activity to increase in a recurrent loop be prevented when there was no earlier motivation? A prominent candidate for a signal that acts after eclosion is juvenile hormone (JH)—though not for reasons that its name implies. Juvenile hormone was first hypothesized (Wigglesworth, 1936) and then shown (reviewed in Riddiford, 2008; Wigglesworth, 1965) to prevent insect larvae from undergoing metamorphosis, and it is largely this larval “juvenile” phase and other developmental functions that have been studied to date (Riddiford, 2008; Robinson, 1992). In *Drosophila*, the predominant juvenile hormone species is methyl 6,7;10,11-bisepoxyfarnesoate (JHB_3_) (Figure 4A) (Richard et al., 1989), and its titer has been reported to increase around the time of eclosion of the imago (Bownes and Rembold, 1987). We verified this earlier report, using liquid chromatography–mass spectrometry, finding a ∼500-fold spike of JHB_3_ titer at eclosion that decays over ∼2 days (Figure 4A).

**Figure 4.**
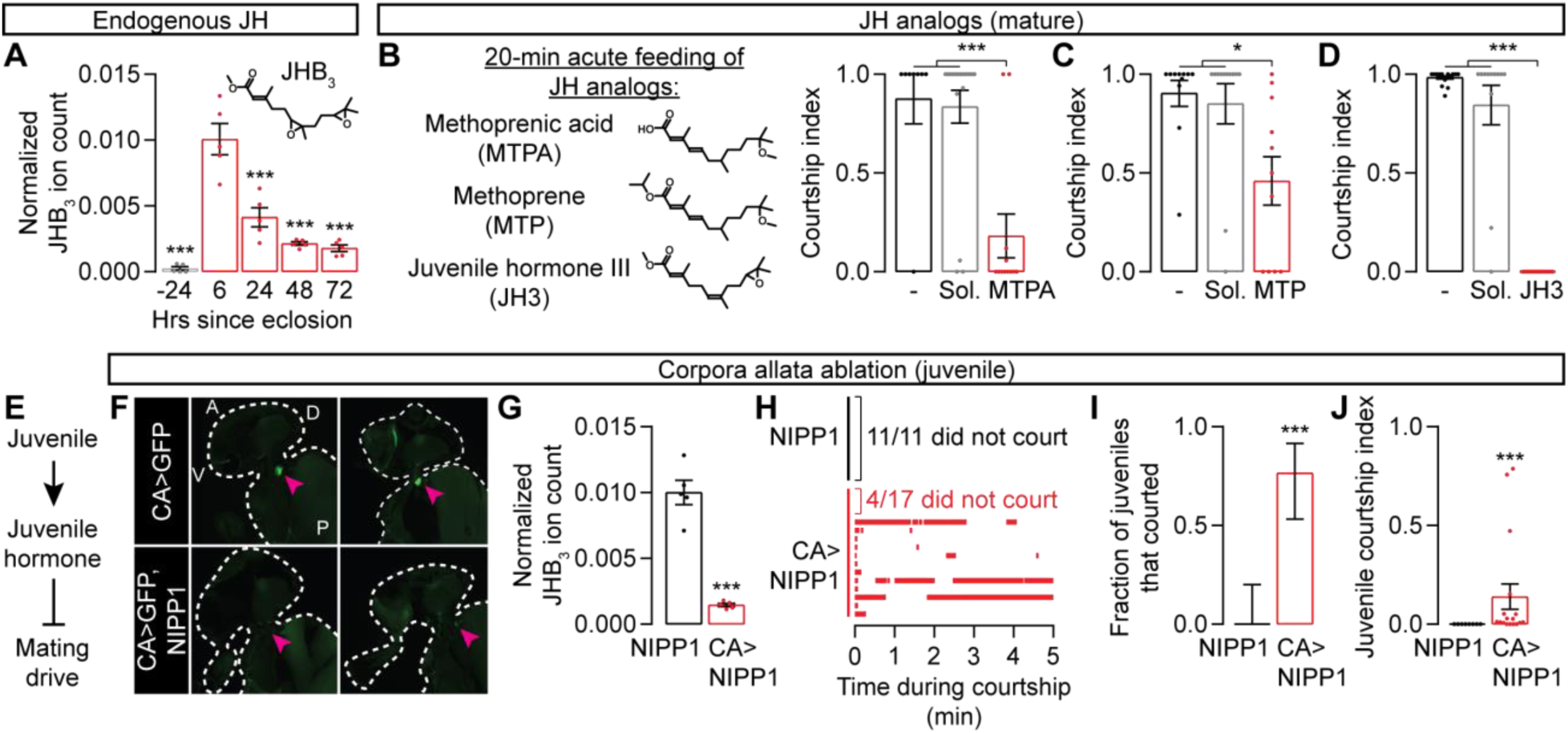
High juvenile hormone titer suppresses mating drive in juvenile males. (A) JHB_3_ titer increases 500-fold around eclosion and decreases over 2-3 days (one-way ANOVA, only comparisons to the 6-hour group are indicated, n = 5 groups of 5 males each). See STAR Methods for quantification details. (B-D) Acutely feeding 3-day-old males with three JHB_3_ homologs decreases mating drive (one-way ANOVA, B: n = 8-19 males, C: n = 11-12, D: n = 12-22). The minus sign indicates flies that were fed neither juvenile hormone analogs nor solvent. (E) Model: high juvenile hormone titer suppresses mating drive in juveniles. (F) Sagittal cryosections of flies (two examples per condition) with or without corpora allata ablation. Arrowheads point to where the corpora allata are—or would be. Outlines trace the fly’s head and upper thorax. Letters denote dorsal-ventral and anterior-posterior axes. (G) Corpora allata ablation decreases JHB_3_ titer in juvenile males (t-test, n = 5 groups of 5 males each). (H-J) Corpora allata ablation increases both the fraction of juveniles that court (I) and their courtship index (J) (I: Fisher’s exact test, J: t-test, n = 11-17 males).

Despite this striking titer profile, the few studies of the effects of juvenile hormone on male courtship behavior have focused on older males and have suggested a modest courtship-promoting role (Argue et al., 2013; Lin et al., 2016; Wijesekera et al., 2016). These previous studies either topically applied juvenile hormone analogs (Argue et al., 2013; Wijesekera et al., 2016) or fed the analogs *ad libitum* (Lin et al., 2016). Instead, we starved mature males for 24-48 hours before transferring them to food containing juvenile hormone analogs for 20 minutes, producing a much higher titer (Figure S3A; see Methods). Acute feeding of three different analogs dramatically lowered mating drive (Figures 4B-4D and S3B). This demotivating effect was not due to either the fasting or the solvents (Figures 4B-4D) and lasted long after the males were returned to normal food (Figure S3C, see Figure 7 for later time points). These results suggest a possible function for juvenile hormone in the suppression of juvenile sexual behavior (Figure 4E).

Juvenile hormone is synthesized in an endocrine organ called the corpus allatum (CA) (Richard et al., 1989). Examining the effects of ablating the two corpora allata of early adults is complicated by the fact that changes in juvenile hormone levels are required for larval molting and metamorphosis. Whereas expressing pro-apoptotic factors in the corpora allata results in larval death (Riddiford et al., 2010 and our observations), the Tatar lab found that targeted expression of Nuclear Inhibitor of Protein Phosphatase type 1 (NIPP1) allowed for normal metamorphosis, but ablated the corpora allata in newly eclosed flies (Parker et al., 2002; Yamamoto et al., 2013). We confirmed this method (Figure 4F) and found that ablating the corpora allata caused a dramatic reduction in juvenile hormone titer (Figure 4G). Though control males never court within the first 6 hours after eclosion, ∼75% of males lacking corpora allata exhibited courtship behaviors in a standard courtship assay (Figures 4H and 4I). The courtship from the allatectomized males was sporadic and fragmented (Figures 4H and 4J), consistent with the higher motivational requirement to maintain courtship bouts (Figures 1B and S1A) (Zhang et al., 2018). Our results below argue that this fragmentation is likely due to the lingering suppression from earlier juvenile hormone production.

### Juvenile hormone directly inhibits multiple motivation-promoting circuit elements

A recent report argued that juvenile hormone increases mating drive in older males by upregulating an olfactory receptor (Or47b) in primary sensory neurons, thereby enhancing pheromone detection (Lin et al., 2016). These conclusions were drawn largely from competition assays between 2- and 7-day old males and were carried out in the dark. When they can see, males that are mutant for Or47b courted well (Wang et al., 2011), a result that we reproduce in Figure S4A. Similarly, in our standard, well-lit, one-male-one-female courtship assays, we see little or no reduction in mature males’ courtship when the neurons expressing Or47b are silenced (Figures S4B-S4C). We do not find any increase in courtship when these sensory neurons are stimulated in juveniles (Figure S4D), nor any influence on recovery from satiety in mature males (Figure S4E). These results join those above and below in arguing that the suppression of sexual behavior in juveniles does not rely on adjustments to primary sensory neurons.

Acutely feeding mature males the juvenile hormone analog MTPA caused severe activity decreases in all three sets of courtship-motivating neurons that normally sustain high calcium levels in the SMPa: the pCd/NPF loop neurons and the downstream dopaminergic neurons (Figures 5A-5C). In contrast, MTPA feeding caused no change in the baseline activity of the circuit elements that are activated by sensory stimuli: the P1 courtship command neurons and the CRNs that are activated by copulation (Figures 5D and 5E). Together with the suppression of courtship behavior (Figures 4B-4D), and the endogenous temporal profile of its titer (Figure 4A), these results indicate that juvenile hormone imposes a juvenile-like state on courtship motivation circuitry (Figure 5F). Also similar to a natural juvenile state (and arguing against non-specific poisoning) stimulating either the dopaminergic neurons, the upstream NPF neurons, or the downstream P1 neurons restored high courtship levels to males fed with the juvenile hormone analog (Figures 5G-5J). As in juvenile males, no rescue of courtship behavior was elicited during acute stimulation of the pCd neurons or from silencing the CRNs (Figures S5A-S5B).

**Figure 5.**
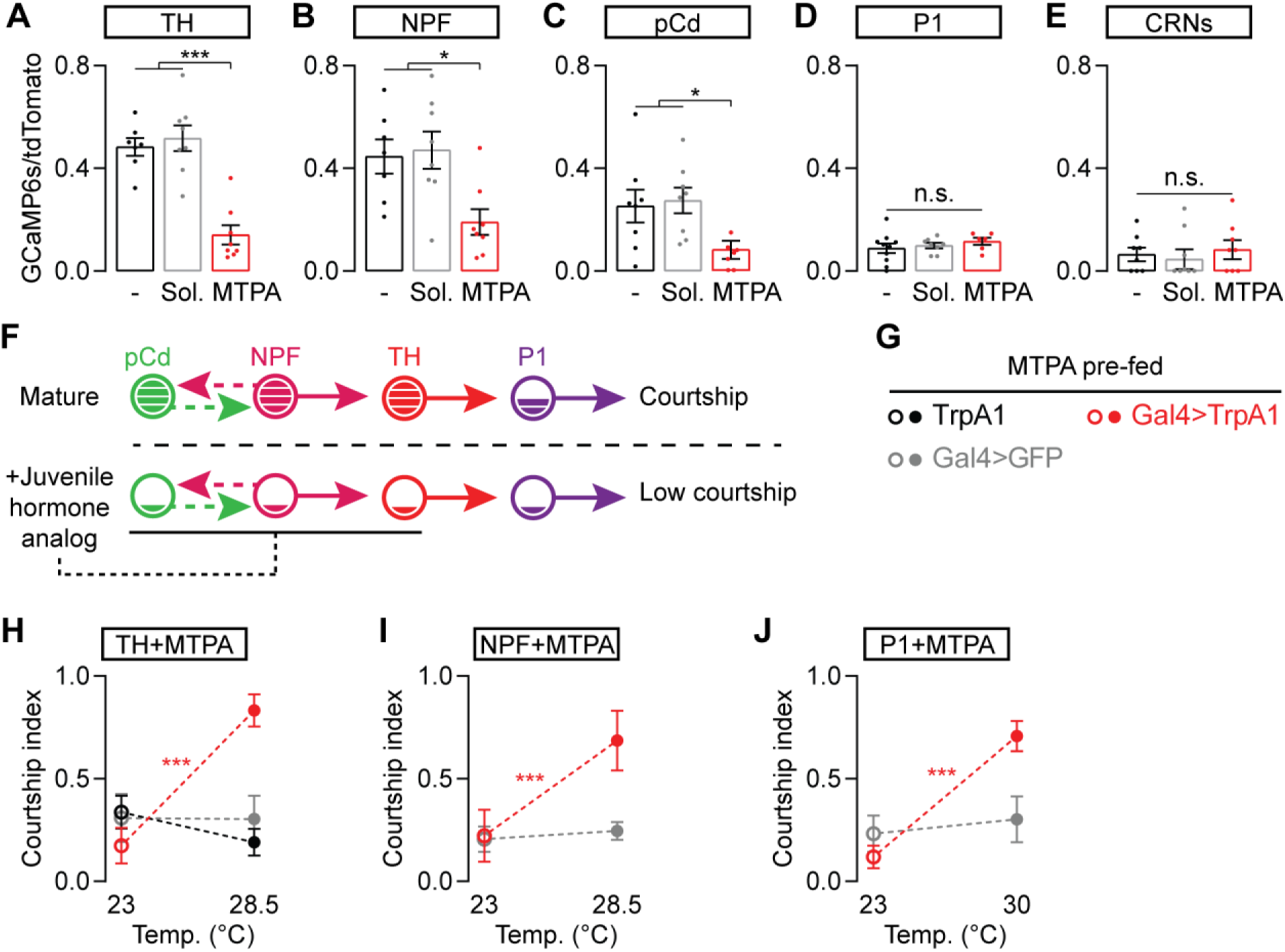
Juvenile hormone signaling suppresses activity in mating-drive circuitry. (A-E) Feeding mature males the juvenile hormone analog MTPA decreases baseline calcium activity in dopaminergic (A), NPF (B), and pCd neurons (C) (one-way ANOVA, A: n = 7-8 male heads, B: n = 7-8, C: n = 7-8, D: n = 6-10, E: n = 8 each). The minus sign indicates the no-food control, which tests the possibility that low courtship is due to the flies not eating the drug-containing food. (F) Model summarizing the results in A-E. (G-J) Acute thermogenetic stimulation of dopaminergic (H), NPF (I), and P1 neurons (J) recovers mating drive after MTPA feeding (two-way ANOVA, H: n = 15-22 males, I: n = 12-16, J: n = 12-21; see Methods for an explanation of stimulation temperatures).

There are two known juvenile hormone receptors in *Drosophila*, the closely related bHLH-PAS transcription factors Methoprene-tolerant (Met) and Germ cell-expressed (Gce). Met and Gce have redundant functions during larval development (Abdou et al., 2011), but separable roles post-metamorphosis (Bilen et al., 2013). As previously reported (Wilson et al., 2003), mature males mutant for Met courted poorly, but we found that Gce mutants showed no courtship defects (Figure 6A). In fact, when we used a tap-induced courtship assay (Zhang et al., 2018), which measures the probability that each tap will lead to courtship and therefore removes the ceiling effect of standard courtship assays, we found that Gce mutants initiated courtship on a higher fraction of taps, indicating hypersexuality (Figure 6B and S5C). Strikingly, feeding juvenile hormone analogs had no effect on either the hypo- or hyper-sexual phenotypes of mutations in the two receptors (Figures 6A-6B and S5C), providing further evidence against poisoning by the drugs and demonstrating that both known receptors are required to receive the juvenile hormone signal. Previous studies have shown that juvenile hormone disrupts Met-Gce dimerization (Godlewski et al., 2006), suggesting a plausible model in which juvenile hormone releases Gce from binding to Met to allow the hormone’s courtship-suppressing functions (Figure 6C). Regardless of the inadequately understood details of its signal transduction, these results show that juvenile hormone works through its known receptors to instruct the mating drive of the male. However, we only rarely observed any courtship from Gce mutant juvenile males (Figure 6D), indicating that juvenile hormone has courtship-suppressing effects that are independent of Gce.

**Figure 6.**
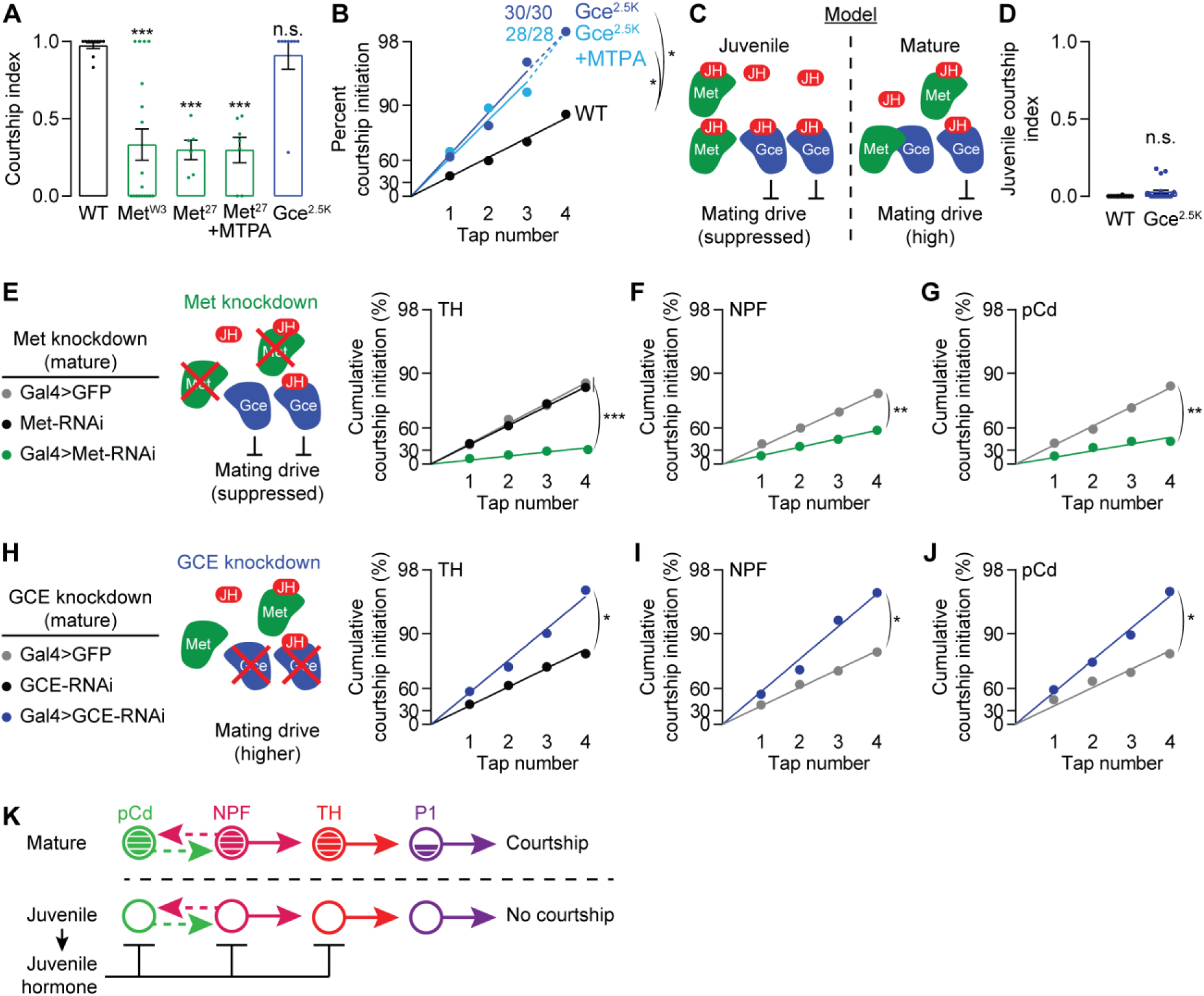
Juvenile hormone directly suppresses multiple components of mating-drive circuitry. (A) Mutation of the juvenile hormone receptor Met decreases mating drive in mature males and renders courtship insensitive to MTPA feeding, while mutation of a second juvenile hormone receptor, Gce, does not affect mating drive in a standard courtship assay (one-way ANOVA, only comparisons to the wild-type flies are indicated = 7-18). (B) Using a tap-induced courtship assay that removes the ceiling effect of standard assays shows that mutating Gce increases mating drive and makes courtship impervious to MTPA feeding (bootstrap, n = 28-32 males) (C) Model: juvenile hormone may disrupt Met-Gce dimerization (Godlewski et al., 2006), allowing Gce to suppress mating drive. (D) Mutation of GCE only rarely causes courtship in juvenile males. (E-G) RNAi knockdown of Met in neurons that store mating-drive information – dopaminergic (E), NPF (F), and pCd neurons (G) – decreases courtship behavior (bootstrap, E: n = 30-39 males, F: n = 28-47, G: n = 29-32). (H-J) Knocking down GCE in dopaminergic (H), NPF (I), and pCd neurons (J) increases mating drive (bootstrap, H: n = 30 males each, I: n =28-33, J: n = 29-30). (K) Model: juvenile hormone directly suppresses the activity of multiple courtship drive-promoting neuronal populations.

**Figure 7.**
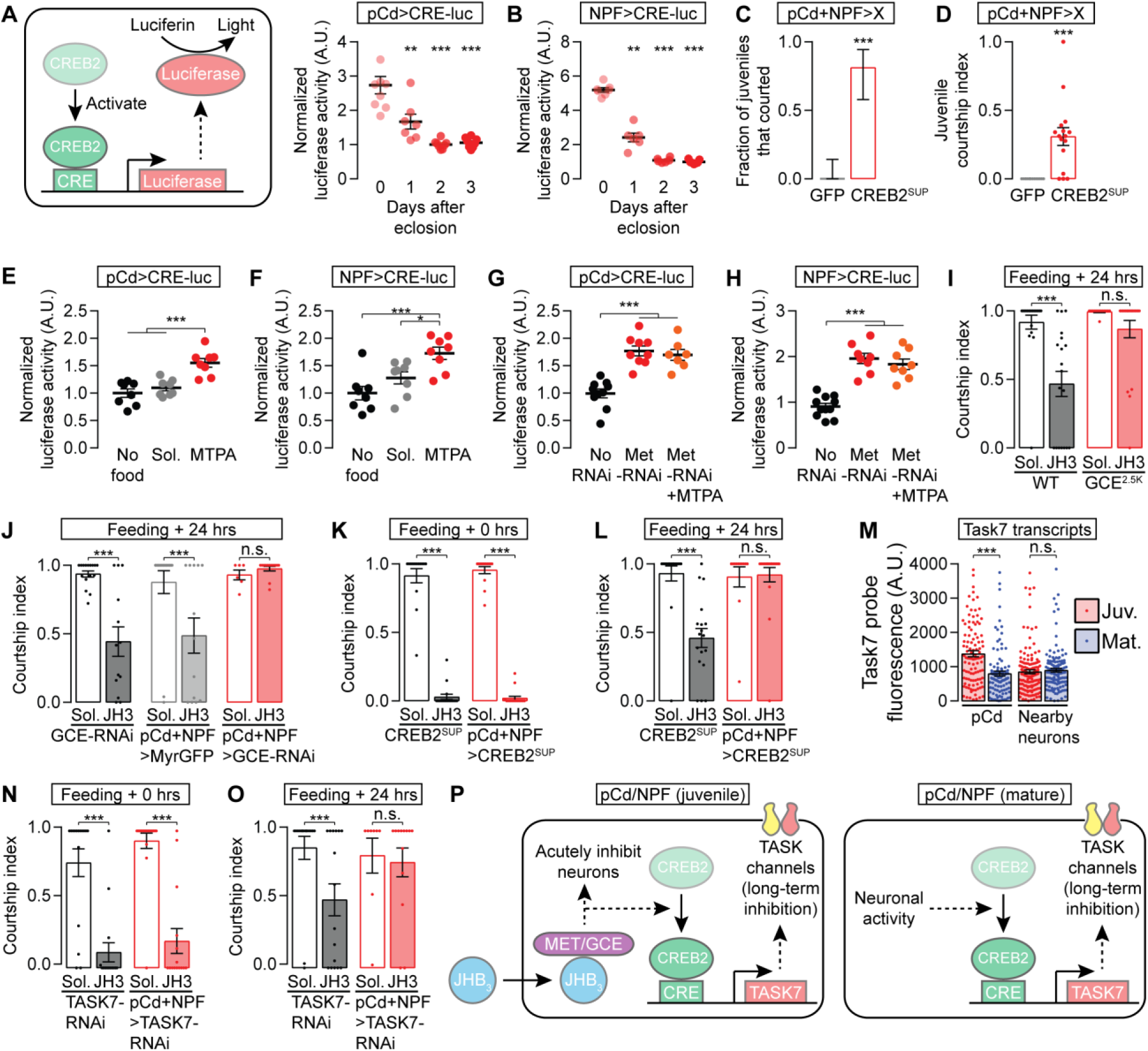
Juvenile hormone causes enduring suppression of mating drive through CREB2-mediated transcription of a potassium channel subunit. (A-B) A luciferase-based assay shows that CREB activity in male pCd (A) and NPF (B) neurons is high just after eclosion and decreases over the ensuing 24-48 hours (one-way ANOVA, only comparisons to the day zero groups are indicated, A: n = 7-8 groups of 3 brains, B: n = 6-7 groups). (C-D) Decreasing CREB2 activity in pCd and NPF loop neurons disinhibits juvenile courtship (C: Fisher’s exact test, D: t-test, n = 16 males each). (E-F) MTPA feeding increases CREB activity in pCd (E) and NPF (F) loop neurons (one-way ANOVA, E: n = 8 groups of 3 brains each, F: n = 8 groups each). (G-H) RNAi knockdown of Met in pCd (G) and NPF neurons (H) both mimics and precludes the CREB-activating effect of juvenile hormone (one-way ANOVA, G: n = 7-9 groups of 3, H: n = 8-11 groups). (I-J) Suppression of courtship lasts at least 24 hours after feeding the juvenile hormone analog JH3, and this effect is blocked by GCE mutation (I) or RNAi knockdown in the pCd/NPF loop (J) (one-way ANOVA, I: n = 13-21 males, J: n = 6-16). (K-L) CREB2 suppression in the pCd/NPF loop does not affect the acute inhibition of courtship by JH3 feeding (K), but prevents the long-term reduction of courtship behavior (L) (one-way ANOVA, K: n = 14-17 males, L: n = 12-20). (M) In pCd neurons (which are identifiable with smFISH by their expression of doublesex transcripts), TASK7 transcripts are expressed at a higher level in juveniles than in adults (one-way ANOVA, n = 105-142 cells). (N-O) TASK7 knockdown in the pCd/NPF loop does not affect the acute inhibition of courtship by JH3 feeding (N), but eliminates the long-term reduction in courtship behavior (O) (one-way ANOVA, N: n = 15-18 males, O: n = 8-16). (P) Model for the fast and slow mechanism through which juvenile hormone suppresses mating drive, compared to satiety-inducing mechanisms in the mature animal.

Though we do not fully understand the mechanisms by which the juvenile hormone signal is transduced to suppress courtship, the involvement of two receptors allowed us to ask whether the hormone acts on previously identified motivational circuitry. We found reductions in courtship when Met was knocked down in the dopaminergic neurons or in either of the populations of loop neurons (Figures 6E-6G and S5D). These effects were relatively modest, consistent with the idea that inhibition at one level of the circuit can be overcome by activity in other motivating neurons. Inversely, courtship probability was increased if any of these courtship-motivating populations were targeted with Gce-RNAi (Figures 6H-6J). No effect was seen when Met or Gce levels were decreased in P1 courtship command neurons or in the CRNs (Figure S5D-S5F). The direct suppression of all known circuit elements that sustain motivational signals by juvenile hormone is in contrast to the action of the CRNs, which, so far as we can tell, rapidly inhibit only the NPF neurons, with consequent and indirect effects on the pCd and dopaminergic neurons (Zhang et al., 2019). The multi-tiered suppression of motivational circuitry seen in juvenile males (Figure 6K) likely explains their complete lack of mating drive, whereas even highly satiated mature males still display some courtship behavior (Zhang et al., 2016).

### Juvenile Hormone activates CREB2 for long-term suppression of mating drive

The juvenile hormone titer that spikes at eclosion decays away by 48 hours (Figure 4A), but mating drive continues to increase for several days (Zhang et al., 2016), suggestive of an enduring courtship-suppressing mechanism. Recently, we found that the satiety state in mature males is maintained and slowly released by CREB-controlled transcription of inhibitory effectors such as the potassium channel subunit TASK7 in the pCd/NPF loop (Zhang et al., 2019). Using a luciferase-based reporter of CREB activity targeted to either the pCd or NPF loop neurons, we found a large spike in CREB activity at eclosion that declined over 48 hours, mirroring the dynamics of juvenile hormone (Figures 7A and 7B). Disrupting CREB activity in the loop by expressing a suppressive isoform of the fly version of CREB (CREB2^SUP^) elicited courtship from juveniles (Figures 7C-7D), demonstrating CREB2’s critical role in the inactivation of courtship circuitry.

In mature males, CREB2 is activated in the pCd/NPF loop by the electrical activity that motivates courtship (Zhang et al., 2019). In juveniles, there is no prior motivation, and the NPF and pCd neurons are inactive (Figures 3B and 3C), suggesting a neuronal activity-independent potentiation of CREB2 transcription. When we fed mature males the juvenile hormone analog MTPA we found increased CREB activity specifically in the pCd and NPF loop neurons (Figures 7E-7F and S5G-S5I). RNAi-mediated knockdown of the juvenile hormone receptor Met in either pCd or NPF neurons elevated CREB activity, and these effects were not enhanced by feeding juvenile hormone (Figures 7G-7H). These results argue that the early spike in juvenile hormone activates CREB2 to impose a long-lasting suppression of loop activity.

The involvement of transcriptional mechanisms in the inactivation of motivational circuitry suggests an enduring effect of the spike in juvenile hormone at eclosion. Late into this study, we were able to obtain juvenile hormone 3 (JH3), structurally the closest and, we find, the most potently demotivating analog of endogenous JHB_3_ (Figure 4D). Feeding mature males JH3 for 20 minutes led to a suppression of mating behaviors that was still evident 24 hours later (Figure 7I), an effect which required the Gce receptor in loop neurons (Figures 7I and 7J). When CREB2 activity was suppressed in the loop, courtship was inhibited immediately after JH3 feeding (Figure 7K), but the long-lasting reduction in courtship was reverted (Figure 7L). We previously found that CREB2 sustains satiety in loop neurons by transcribing the leak potassium channel subunit TASK7 (Zhang et al., 2019). Similarly, we find that the transcription of TASK7 (quantified using smFISH) is higher in the pCd neurons (which are identifiable by their location and the expression of doublesex) of juveniles than of mature males (Figure 7M). As expected from a gene product that must be transcribed, translated, and trafficked to the membrane to be effective, knocking down TASK7 in the loop had no effect on the immediate inhibition of courtship in mature males but, like CREB2 activity suppression, prevented the enduring effects of JH3 feeding (Figures 7N and 7O). These results underscore the multi-tiered nature of juvenile hormone’s suppression of motivation: at the circuit level (direct action on dopaminergic, pCd, and NPF neurons), over time (immediate and long lasting), and at the molecular level (CREB2 and TASK7 for long-lasting suppression in the loop, but other unidentified mechanisms for immediate suppression and in the dopaminergic neurons) (Figure 7P). These mechanisms ensure the complete suppression of sexual motivation, which, propelled by the recurrent excitation loop, has a natural tendency to increase over time.

## Discussion

“The snake in the story of Eden is doubtless a phallic symbol…suggesting sexual awakening as the beginning of the knowledge of good and evil” (Durant, 1954). The transition to adulthood has profound consequences for many brain functions, as was appreciated long before those functions were known to derive from the brain. The hormonal signaling pathways that drive these changes in mammals are now understood, but comparatively little is known about their specific effects on behavioral and motivational circuitry. Teleological explanations for delays in the onset of sexual behavior may vary across species, but the pervasiveness of this phenomenon suggests the possibility of a core mechanistic conservation that transcends idiosyncrasies of duration, purpose, and the molecular nature of the hormonal trigger. The system we develop here for studying the transition to sexuality allows rapid insight, with principles that suggest molecular and circuit hypotheses in other animals and other late-emerging brain functions.

One obvious, but likely superficial difference in the control of behavioral maturation between *Drosophila* and mammals is the sign of the regulation by circulating hormones. Though some gonadal hormones can suppress mating drive (Freund, 1980), in juvenile male rodents, experimental elevation of testosterone causes precocious mating behavior (Södersten et al., 1977), whereas we find that juvenile hormone suppresses mating behavior. This distinction becomes less prominent early in signal transduction, as loss of the Met receptor for juvenile hormone decreases sexual behavior, similar to the effects of androgen (testosterone) receptor antagonists in humans (Boccardo et al., 1999). The receptors for testosterone, estrogen, and progesterone are all transcription factors, as are the Met and Gce receptors for juvenile hormone. Decades of work on juvenile hormone signaling during the larval and pupal phases of *Drosophila* life have identified the two receptors that have guided our studies here, but, unlike ablation of the source of juvenile hormone or overriding its action at the level of neuronal activity or CREB activation, removing the suppressive receptor, Gce, does not cause substantial juvenile courtship. This points to the likely existence of yet another juvenile hormone receptor—or a second signaling system—the identification of which will provide a deeper understanding of the suppressive mechanisms used to completely, selectively, and transiently inactivate this fundamental drive.

In mature animals, the recurrent loop that drives the male to court also primes itself for satiety through activity-induced, CREB-mediated production of suppressive TASK7-containig potassium channel complexes (Zhang et al., 2019). This motivation-suppressing module is also used in juveniles, with hormonal signaling causing CREB2 activation. Though we have not yet identified additional suppressive effectors in juveniles, we see evidence of fast-acting suppression by juvenile hormone, as well as direct suppression of the dopaminergic neurons, neither of which can be accounted for by CREB activity in the pCd/NPF loop. Parallel fast and slow responses have also been found in the response to mammalian gonadal hormones (e.g., Levin, 2009). Multiple repressive mechanisms are likely necessary to completely inactivate mating drive, since individually removing or overriding individual suppressive effectors allows juvenile courtship. This is again similar to findings in mammals, where administering testosterone to either the preoptic area of the hypothalamus or to the medial amygdala suffices to restore mating behaviors in castrated males (Coolen and Wood, 1998; Putnam et al., 2001).

A second clear difference between *Drosophila* and vertebrate juvenile stages is the timescale. The fly system requires suppression of sexual behavior for days, not months or years (even 150 years as suggested in Greenland sharks (Nielsen et al., 2016)). But several non-neuronal structures are still maturing in juvenile flies, such as the abdominal musculature, the cuticle, and the ejaculatory bulb, leading to the *a priori* expectation that the neuronal circuitry that will eventually animate these features may require structural development as well. Though we have not undertaken the kind of detailed anatomical analysis required to argue for or against fine-scale structural rearrangements, courtship circuitry appears grossly normal in juveniles, and stimulation of several circuit elements (P1, Dopaminergic, NPF, and pCd) can rapidly drive coherent courtship. These findings serve as strong evidence against a strict developmental ontogeny for the post-eclosion emergence of reproductive motivation and behavior. This appears at odds with the recent report of structural changes in a key hypothalamic population during estrus, and with the inability of these neurons to drive sexual behaviors when activated in ovariectomized mice (Inoue et al., 2019). We note that we are not able to restore sexual behavior in male flies if we stimulate only subpopulations of individually necessary dopaminergic neurons (Figure S2B) (Zhang et al., 2016), a consideration that, we argue, leaves open the possibility that the motivational and behavioral circuitry in female mice may be at least partially functional, but suppressed, in non-estrus mice. The most parsimonious explanation may be that both structural and activational changes take place between asexual and sexual periods of life. While the findings of the mouse estrus study (Inoue et al., 2019) advance the hypothesis that the structural rearrangements drive the changes in circuit activity, our results suggest it may be the other way around: increased functional connectivity could result from activity-dependent mechanisms, as has been reported in a growing number of systems (Liu et al., 2017; Yan et al., 2017). We can find no reports of synthetic stimulation producing juvenile sexual behavior in the mammalian literature, but stimulation of hypothalamic neurons can produce coherent and robust parenting behavior from otherwise non-parental mice (Wu et al., 2014), demonstrating latent but functional circuitry for these late-emerging behaviors.

Finally, our results provide an alternative explanation for what has been the strongest evidence in favor of the organizational hypothesis for the maturation of reproductive behaviors in mammals: the delay between hormonal changes and the behaviors they induce (Schulz et al., 2009). In the fly, the long-lasting, CREB-imposed effects of an earlier suppressive hormonal state must decay away to allow loop activity to ramp up and promote sexual behavior. Enduring and distributed suppressive effects may explain why the awakening of sexuality triggered by hormonal changes is gradual and halting in both flies and mammals.

## Acknowledgements

We thank the Rogulja and Crickmore labs for comments on the manuscript. Mark Andermann and Chuck Weitz provided equipment for two-photon microscopy and luciferase assays, respectively. Lisa Goodrich and Michelle Frank provided equipment and assistance for whole-fly cryosectioning. Mass spectrometry was conducted at the Harvard Center for Mass Spectrometry. Michelle Frank, Christine Boutros, Lauren Miner, Ben Gorko, Ariadna Corredera, and Blake Karavas helped with experiments. S.X.Z., E.G., and M.A.C. performed the experiments. All authors designed the experiments, analyzed data, and wrote the manuscript. S.X.Z. is a Stuart H.Q. & Victoria Quan Fellow at Harvard Medical School. D.R. is a New York Stem Cell Foundation-Robertson Investigator. This work was supported by The New York Stem Cell Foundation and a grant from the NIH (DP1DA044358). The authors declare no conflict of interest.

## STAR Methods

### CONTACT FOR REAGENT AND RESOURCE SHARING

Please contact D.R. or M.A.C. for fly strains and other resources in this study.

### EXPERIMENTAL MODEL AND SUBJECT DETAILS

#### Fly Stocks

Fly husbandry was performed as previously described (Zhang et al., 2016). Flies were maintained on conventional cornmeal-agar medium under a 12-hr light/12-hr dark cycle at 25 °C and ambient humidity. Due to the focus of this study, all tested flies were adult males. Unless otherwise stated, males were collected on the day of eclosion and group-housed away from females for either 3-6 hours (juvenile) or 3-4 days (mature) before testing. Virgin females were generated using a hs-hid transgene integrated on the Y-chromosome (Boutros et al., 2017; Dietzl et al., 2007). All behavioral experiments were carried out within the first 10 hours of each light day-phase. Detailed genotypes of all strains used in the paper are listed in Table S1 available online.

### METHOD DETAILS

#### Courtship Assays

Courtship assays were used to test if a manipulation decreases mating drive and were carried out in the same setting as previously described (Boutros et al., 2017; Zhang et al., 2016). A male fly (3-4 days old) and a w^1118^ virgin female fly were videotaped in cylindrical courtship chambers (10 mm diameter and 3 mm height) at 23 °C and ambient humidity. In experiments where we looked for both potential increases and decreases in courtship, we conducted experiments analyzing tap-induced courtship initiations, a metric that is less likely to reach ceiling than the standard courtship index (Zhang et al., 2018). These experiments are conducted in the same way as the regular courtship assays, but recorded with a more zoomed-in camera and analyzed differently (see below). In Figures S1B-S1C, we measured courtship by mature males that are sexually satiated. In those experiments, each male was paired with 15 virgin females for 4.5 hours to ensure satiety (Zhang et al., 2016), and then tested after 0-3 days of being away from females (recovery).

For thermogenetic experiments using TrpA1, half of the males were assayed at 23 °C, and the other half at a higher temperature (28.5 °C or 30 °C). We typically started testing with 30 °C, and if the flies show abnormal behaviors (e.g., seizure in TH>TrpA1 flies), we lower the temperature to 28.5 °C. We previously showed that, in mature satiated males, acute dopaminergic stimulation instantly restores robust courtship initiations, but it requires several minutes of dopaminergic stimulation for the flies to sustain courtship bouts (Zhang et al., 2018). Similar observations are also seen here in juvenile males (Figures 2F). To ensure the consistency of stimulation, unless otherwise stated, stimulation of the dopaminergic and its upstream neurons are preceded by a 20-minute pre-stimulation at the same temperature

For conditional silencing experiments involving TubGal80^ts^ (e.g., Figure 5L), male flies were moved to 30 °C after eclosion and kept there (isolated from females) until the assay, which took place at 23 °C.

For the abdomen-removal experiment (Figure S1D), the flies were anesthetized on ice, and their abdomens were surgically removed as described in (Zhang et al., 2016).

#### Satiety Assay and Reversal

Satiety assays were used to test if a neuronal population (e.g., Or47b neurons) has an acute motivation-promoting role and were carried out as previously described, with minor modifications (Zhang et al., 2016). Individual male flies were placed with ∼15 virgin females in a food vial at 23 °C and ambient humidity for 4.5 hours. Mating behaviors (courtship and copulation) were scored manually at time points: 0.5, 1, 4 and 4.5 hours. 4.5-hour satiety assays were used to satiate males for courtship and imaging experiments.

To thermogenetically revert satiety, the temperature was raised after the last time point, and mating behaviors were scored 20 minutes after the incubator reached the appropriate temperature.

#### Quantification of Endogenous JHB_3_

To quantify JHB_3_, groups of 5 flies were homogenized and then mixed with 100 μL methanol to extract the hormone. The mixture was spun down at 17000 × g for 20 seconds, and the supernatant was transferred to a new vial and diluted 3 times with dH_2_O (1 part supernatant and 2 parts dH2O). 20 μM of exogenous Juvenile Hormone 3 was added to the solution as an internal standard for quantification (controlling for the deviations of the instrument performance from experiment to experiment). If needed, samples were stored at −20 °C prior to quantification.

We used liquid chromatography coupled with mass spectrometry to quantify JHB_3_. Each sample was first loaded in a standard reverse-phase C18 column (Agilent, ZORBAX SB-C18) by a high-performance liquid chromatography (Agilent 1200 Series). Each sample was then eluted off the column over the course of 23 minutes using two solvents, ddH_2_O (with 0.1% formic acid) and acetonitrile (with 0.1% formic acid), and analyzed with a mass spectrometer (Agilent 6210 Time-of-Flight mass spectrometer) in cation mode. During each elution, acetonitrile concentration was ramped up from 5% (T = 2 min) to 100% (T = 13 min), kept at 100% for 5 min, and then ramped down from 100% (T = 18 min) to 5% again (T = 18.1 min). JHB_3_ and JH3 were consistently detected at T = ∼10 min and T = ∼12 min respectively.

#### Acute Feeding of Juvenile Hormone Analogs

To ensure consistent drug feeding (and juvenile hormone titer), male flies were starved (but not water deprived) in vials containing 5 mL of 0.8% agarose gel (made with dH_2_O) for 36 hours prior to feeding. However, for less starvation-resistant genotypes (>10% lethality at the end of starvation), the time was shortened to 24 hours. The exact time of starvation was kept consistent within each genotype. For feeding, flies were transferred to vials containing 2 mL of 0.8% agarose gel (made with 50% dH2O and 50% grape juice) with a juvenile hormone analog for either 20 minutes (if the flies were tested right away) or 3 hours (if the flies were tested 24 hours later). To assist feeding, the vials are placed upside down so that the flies can easily find the food anti-geotactically. The concentrations of the drugs were 200 μg/mL for methoprenic acid (MTPA), 200 μg/mL for methoprene (MTP), and 168 μg/mL Juvenile Hormone III (JH3).

#### Whole-fly Cryosection and Imaging

Whole flies with GFP-expressing corpora allata were briefly dipped in 75% ethanol to remove cuticle hydrocarbons before being fixed in 4% paraformaldehyde dissolved in PBS with 0.3% Triton X-100 (PBST) for 60 minutes. After 3x 20-minute washes with PBST, flies where transferred into Neg-50 medium (ThermoFisher, 6502) and frozen in a dry ice/acetone cooling bath (−78 °C). The frozen flies were sliced sagittally into 100-μm sections in a cryosect and mounted directly on a glass slide with mounting medium (NEG-50). Confocal sections (4-7 sections at 3-µm intervals) were taken from the medial sections where corpora allata are seen (if not ablated).

#### Antibody Staining and Two-photon Volume Microscopy

Antibody staining was performed as described before (Zhang et al., 2016). Myristoylated GFP, which was driven with different Gal4 lines, was stained with chicken anti-GFP (1:1000) and then donkey anti-chicken (1:400) antibodies. Two-photon microscopy was performed using a Neurolabware microscope and a tunable pulse laser (InSight X3, Spectra-Physics) as described before (Burgess et al., 2016). Volume imaging was done with an 8 kHz resonant scanning mirror (CRS8, Cambridge Technology) along the x-axis, a galvanometer scanning mirror (6215H, Cambridge Technology) along the y-axis, and an electrically tunable lens (EL-10-30-TC-NIR-12D, Optotune) along the z-axis. For each sample, we imaged 20 volumes (31 frames per volume and 15.5 frames per second spanning 100 μm along the z-axis) at 960 nm, calculated a median volume to cancel out PMT-noise, and finally acquired a standard-deviation projection of the volume to visualize the processes.

#### Baseline Calcium Imaging of Dissected Brains

Fly brains expressing GCaMP6s (Chen et al., 2013) and tdTomato were dissected from either juvenile or mature males in HL3.1 solution (Feng et al., 2004) and mounted anterior-side down onto the base of a glass-bottom Petri dish (MatTek, P35G-1.5-20-c) housing 3 mL HL3.1 saline, as previously described (Zhang et al., 2016). Confocal sections (4-7 sections at 3-µm intervals) were taken from the anterior of the superior medial protocerebrum (SMPa). Whenever possible, we also co-expressed and imaged tdTomato in the same cells, to normalize the GCaMP6s signal. Whenever possible, we also imaged from unrelated brain regions (the mushroom body) to show that the changes are specific to the SMPa.

#### Measuring CREB2 Activity

To measure CREB2 activity we used a FLP-dependent luciferase construct driven by the cAMP-response element (CRE) promoter (Tanenhaus et al., 2012), with UAS-FLP driven in the neurons under examination. The luciferase assays were performed using a commercial kit (Promega, E1910). Brains of males were dissected in HL3.1 solution and transferred, in groups of 3, into 50 µL dissociation solution for 5 minutes at −20 °C. Then, a 20 µL supernatant of the dissociation solution was added to 50 µL substrate solution, and the luciferase activity was measured in a photoluminometer (Turner Designs TD-20/20) over a 5-second window.

#### Fluorescent *in situ* hybridization

Whole-mount in situ hybridization was performed as described previously (Zhang et al., 2019). Brains dissected from juvenile and mature wild-type Canton-S males were used for these experiments. Single-molecule fluorescent *in situ* hybridization (smFISH) was performed following the published protocol (Long et al., 2017), with the exceptions that we omitted 1) the overnight incubation in 100% ethanol and 2) the bleaching steps. Fluorescence-tagged TASK7 (Quasar 570) and Dsx (Quasar 670) probes were designed and manufactured using a commercial source (LGC Biosearch Technologies). Confocal sections were acquired using an Olympus Fluoview 1000 microscope at 3-µm intervals.

#### Pharmacology

Methoprenic acid (MTPA, Sigmal-Aldrich, M6682) was dissolved in acetone at 5 mg/mL (18.6 mM). Methoprene (MTP, Sigma-Aldrich, 33375) was dissolved in DMSO at 100 mg/mL (322.1 mM). Juvenile hormone III (JH3, Sigma-Aldrich, J2000, Lot purity = 79%) was dissolved in DMSO at 4.21 mg/mL (15.8 mM).

### QUANTIFICATION AND STATISTICAL ANALYSIS

#### Analysis of Courtship Behavior

The courtship index was manually scored as previously described (Boutros et al., 2017; Zhang et al., 2016). It is defined as the fraction of time during which the male fly is engaged in mating behaviors (courtship and copulation) over 5 minutes after courtship initiation. A bout of courtship was scored as initiated when the male oriented toward the female, began tracking her, and unilaterally extended his wing to sing to her. A bout was scored as terminated if the male stopped tracking the female or turned away from her. Once courtship was terminated, it was almost never reinitiated within 10 seconds. Whenever possible, terminations were verified by confirming that the male did not resume singing when the female subsequently passed in front of him.

For experiments that measure tap-induced courtship probabilities, the experimenter who scored the videos was blind to the genotypes and experimental conditions. Taps are defined as a male foreleg touching any part of the female body. In our assays, courtship is always initiated by a tap, always within a few seconds, and most often within 500 ms. The details of this analysis can be found in Zhang et al (Zhang et al., 2018). Briefly, we calculated the cumulative courtship probabilities (y) after each tap (x). We plotted –Ln(1 – y) against x fitted a straight line that is forced through the point of origin ([0, 0]). The Y-axis ticks are labeled with the same log transformation. This fitted line represents the expected cumulative courtship-initiation probability if the tap-to-courtship transitions are modeled as a perfect coin (with varying odds). A steeper line represents higher courtship probability (i.e., high odds of the coin). From the slope of the line (s), we can calculate the per-tap courtship probability as 1 – e^−s^.

We used bootstrapping to perform hypothesis tests on courtship probabilities, with the underlying null hypothesis that these two datasets were generated with the same unknown courtship probability. Before bootstrapping, we first pooled two datasets together. Then, we resampled two bootstrapped datasets from the pooled data with the same sample size as in the experiments.

Then, using the same linearization and linear regression procedures, we calculated new, bootstrapped courtship probabilities. We calculated their difference and repeated the resampling and re-calculations 100,000 times. The p-value was then calculated as the fraction of iterations that generate a difference in courtship probabilities at least as large as the experimental one. We used Bonferroni corrections to adjust the p-values for multiple comparisons. The MATLAB code to perform these calculations is available online.

#### Analysis of Satiety Assays

For standard 4.5-hr satiety assays, individual males were paired with ∼15 virgin females and mating behaviors (courtship and copulation) were scored manually at time points: 0.5, 1, 4 and 4.5 hours. 4.5-hour satiety assays were used to satiate males for courtship and imaging experiments. A within-group (5-7 males per group) average was generated for each time point. The first two (generally 0.5 and 1 hour) and last two (generally 4 and 4.5 hours) time points were averaged to generate Initial and End percentages, respectively. For satiety reversal, another percentage is generated from the mating and courtship behaviors during stimulation (20 minutes after temperature equilibration).

#### Quantifying Imaging Data

For quantification of baseline calcium level, after one person (S.X.Z.) took the images, another person (D.R. or E.G.) selected regions of interest (ROIs) of ipsilateral SMPa, mushroom body medial lobe, and background (near the antennal lobe, where dopamine projections are sparse) using the tdTomato channel. ROI sizes were kept roughly the same between samples. Average pixel fluorescence in each ROI was calculated with a custom-written MATLAB script. Whenever possible, normalized fluorescence was calculated as (GCaMP6s_SMPa – GCaMP6s_background) / (tdTomato_SMPa – tdTomato_background). The same formulas were used for the mushroom body medial lobe as well. The MATLAB code to perform these step is available online.

For smFISH quantification, we used a custom, semi-automated MATLAB script to segment images and measure fluorescence intensities as described previously (Zhang et al., 2019). Cell bodies were first segmented using the Dsx channel and the TASK7 channel. The TASK7 puncta were then grouped into either a Dsx-positive or a Dsx-negative population, and the average fluorescence intensities were calculated separately. Background fluorescence was measured from an unrelated brain region (the antenna lobe) and subtracted from the data.

#### Analysis of JHB_3_ titer

We extracted the ion counts corresponding to the m/z ratios of monoprotonated JHB_3_ ([M+H]^+^ = 283.1909±0.0057) and the internal control Juvenile Hormone III ([M+H]^+^ = 267.1960±0.0053) from the chromatograph. Peaks corresponding to JHB_3_ and Juvenile Hormone III are consistently seen at T = ∼10 min and T = ∼12 min respectively and integrated to estimate the ion counts. The ion counts of Juvenile Hormone III are reasonable consistent between experiments (standard deviation/mean = 11%). We reasoned that this deviation is largely due to fluctuations of instrumental performances and normalized the ion counts of JHB_3_ (target of quantification) by taking their ratios over the ion counts of the internal standard.

#### Additional Statistical Tests

One-way ANOVA, two-way ANOVA, two-tailed student’s t-test, and Fisher’s exact test were performed using Prism 7. All tests are unpaired. All One-way ANOVA tests use post-hoc Tukey corrections. All Two-way ANOVA tests use post-hoc Bonferroni corrections. Error bars represent 1 S.E.M. for most panels and Jeffreys’ 95% confidence interval for proportions.

#### DATA AND SOFTWARE AVAILABILITY

MATLAB scripts and functions are available online at https://github.com/CrickmoreRoguljaLabs.

---

**Figure S1.**
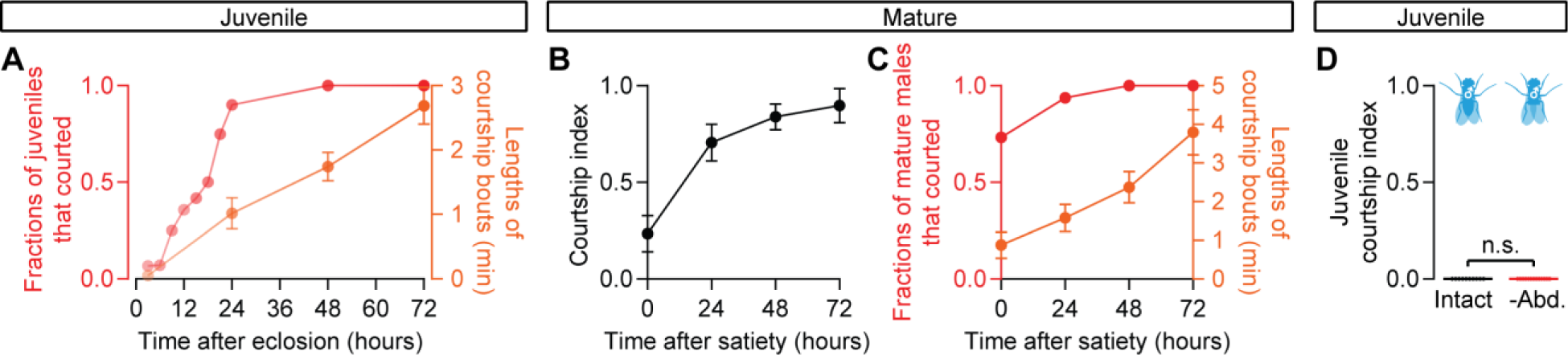
Male mating drive gradually accumulates over the first 3 days after eclosion. (A) Most males have begun to display courtship behaviors by 24 hours of eclosion; their initial courtship bouts are fragmented and gradually become consolidated over the first 3 days (n = 12-32 males). (B-C) Mature males gradually regain courtship vigor after satiety (n = 11-18 males). (D) The lack of courtship in juvenile males is not due a suppressive effect from the reproductive organs, as their surgical removal does not cause courtship (n = 10-13 males).

**Figure S2.**
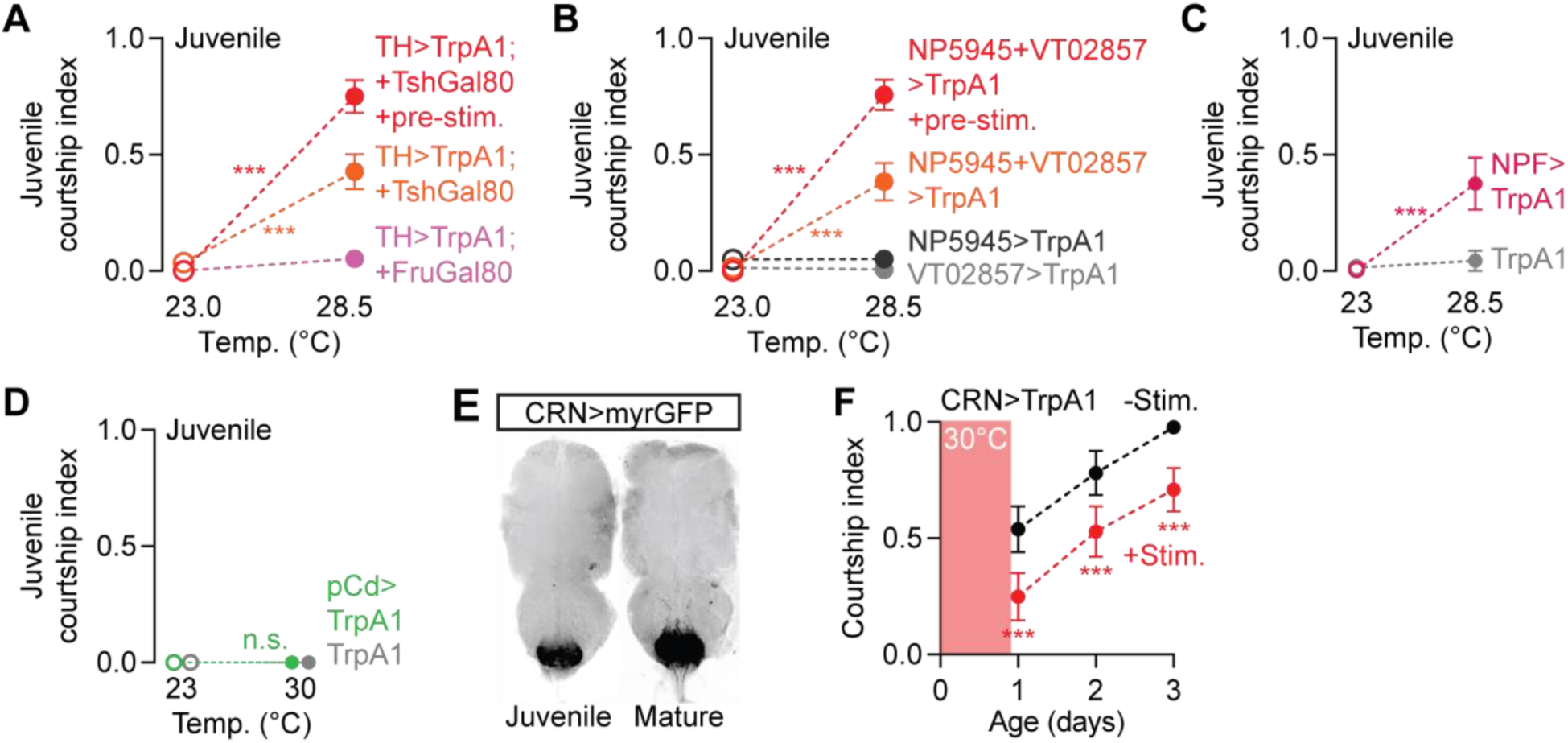
Stimulation of mating-drive circuitry induces courtship in juvenile males. (A-B) Thermogenetic stimulation of all brain dopamine neurons (A) or the mating-drive-promoting subsets (B) induces more vigorous juvenile courtship if the neurons have been pre-stimulated for 5 minutes before the assay. The motivational effects are blocked by a Gal80 transgene indirectly controlled by the Fruitless locus (***p<0.001, two-way ANOVA, A: n = 14-15 males, B: n = 8-16). (C-D) Acute stimulation of NPF (C) but not pCd neurons (D) induces courtship in juveniles (***p<0.001, n.s. not significant, two-way ANOVA, n = 12-16 males, n = 10-15). (E-F) Thermogenetic stimulation of the CRNs delays the onset of courtship behavior (***p<0.001, two-way ANOVA, n = 16-23 males).

**Figure S3.**
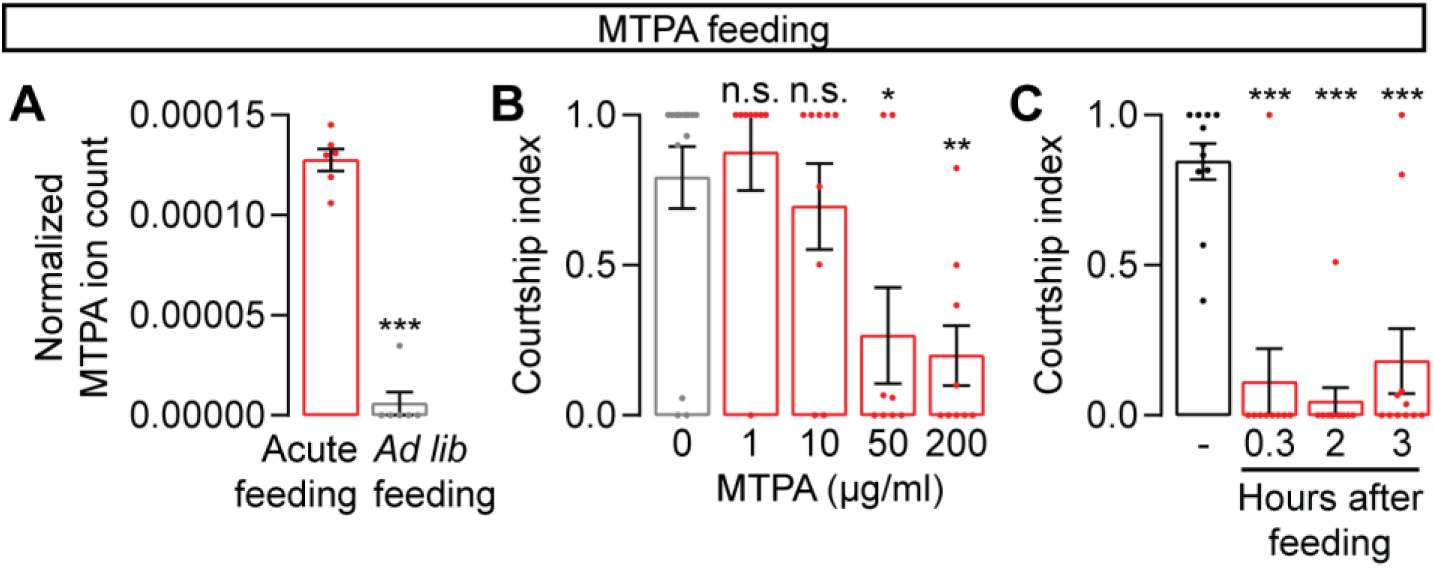
Acute feeding of MTPA decreases courtship in male flies. (A) MTPA titer is much higher in food-restricted males that are fed with MTPA-containing food than in males that are raised on the same food *ad libitum* for 3 days (n = 6 males each). (B) MTPA is effective in decreasing courtship at a concentration of 50 μg/mL (**p<0.01, *p<0.05, n.s. not significant, one-way ANOVA, n = 8-15 males). (C) MTPA suppresses mating drive hours after the 20 minute acute feeding period has ended (one-way ANOVA, n = 8-11 males).

**Figure S4.**
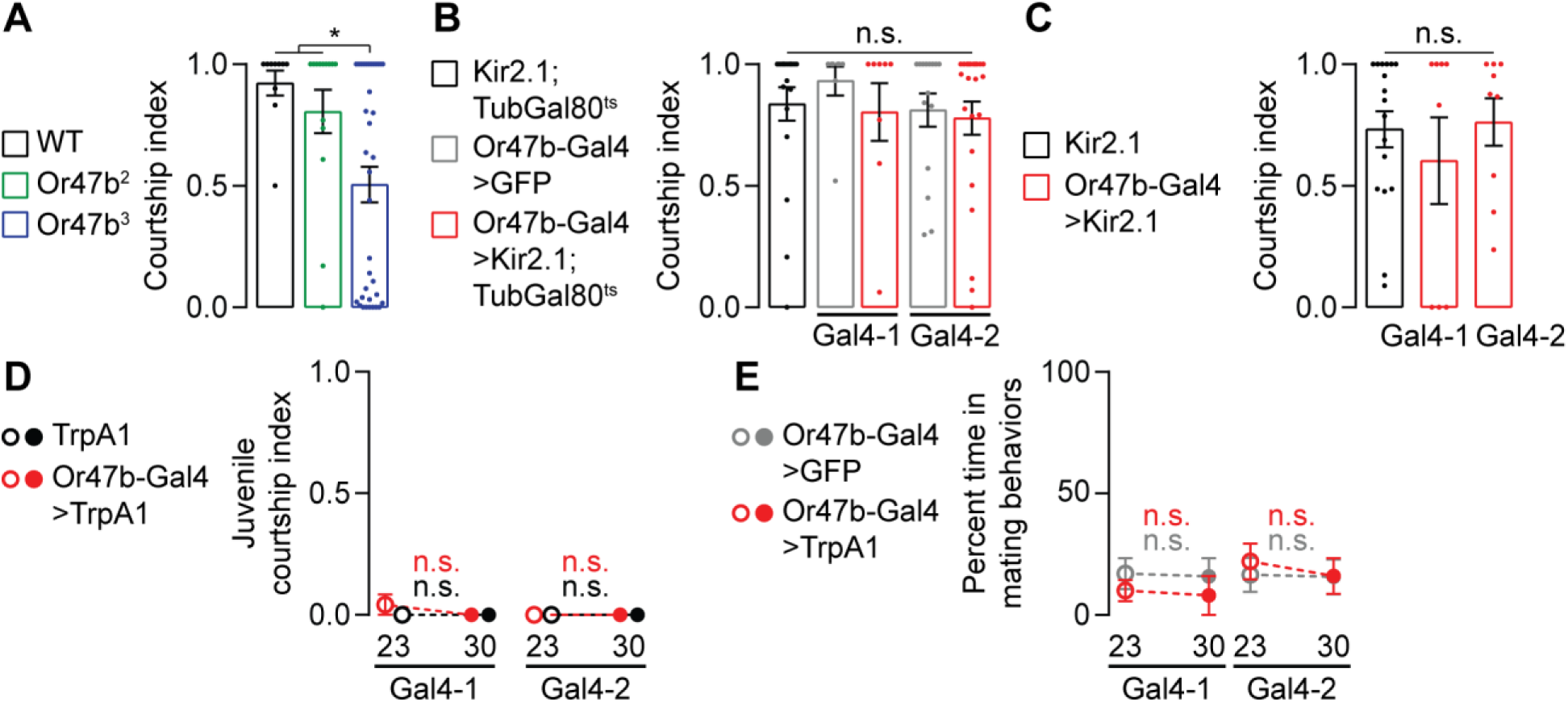
Or47b is not required for courtship produced by 3-day old males in the light. (A) 3-day old male flies mutant for the Or47b null allele (Or47b^2^) used in (Lin et al., 2016) show little, if any, decrease in courtship under well-lit conditions. A different mutant line did show reduced courtship, but we did not further assess the overall health of this stock so the reduction in courtship may be due to background effects (*p<0.05, t-test, n = 10-14 males). (B-C) Silencing Or47b-expressing neurons either conditionally (B) or constitutively (C) does not significantly decrease courtship in 3-day old males (n.s. not significant, one-way ANOVA, B: n = 8-36 males, C: 8-17). (D-E) Thermogenetic stimulation of the Or47b-expressing neurons does not induce courtship in juveniles (D) or revert satiety in mature males (E) (n.s. not significant, two-way ANOVA, D: n = 8-13 males, E: n = 5 groups of 5-7 males each).

**Figure S5.**
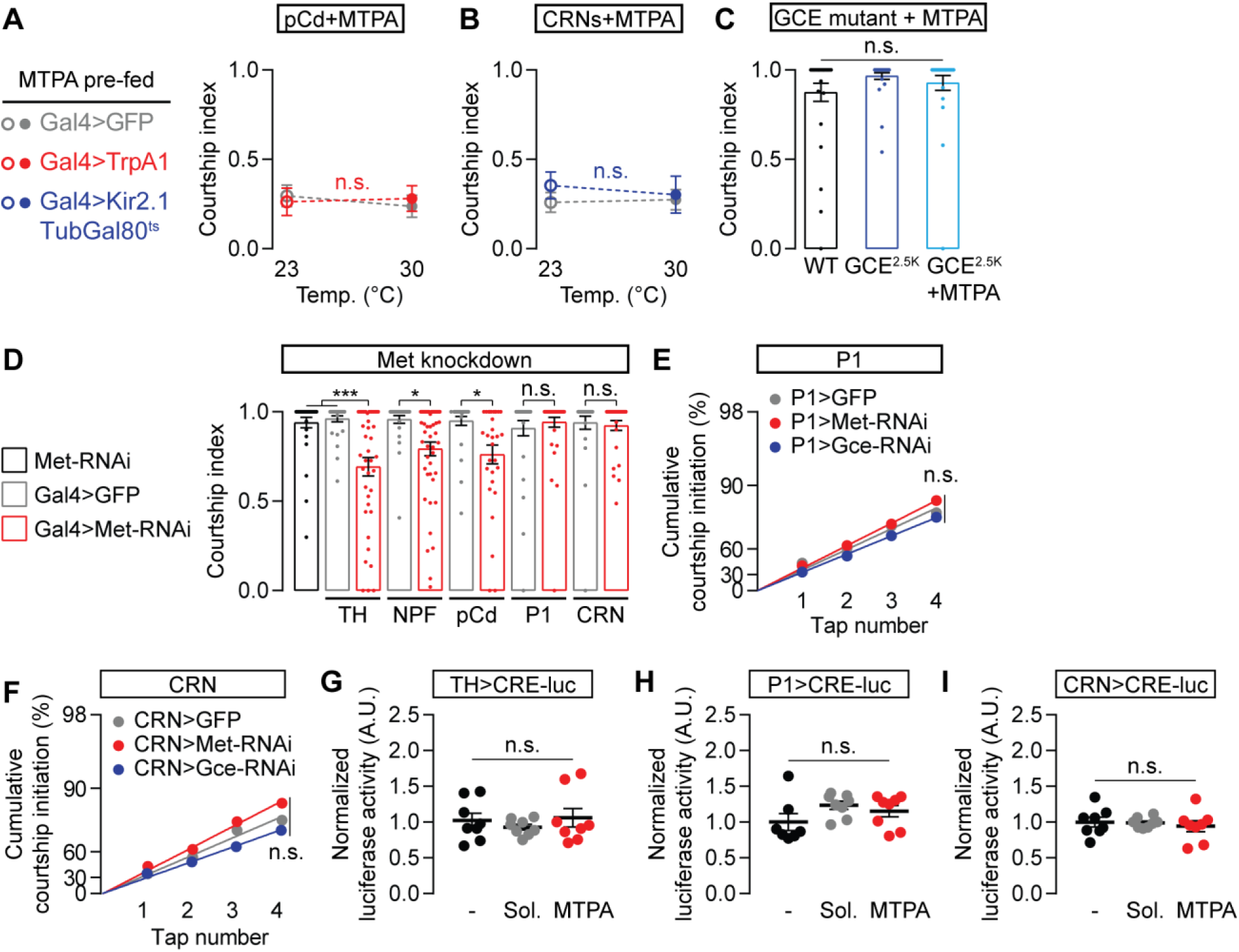
Effects of juvenile hormone signaling on mating-drive circuitry and courtship behavior. (A) Acute thermogenetic stimulation of pCd neurons does not recover mating drive after MTPA feeding (two-way ANOVA, n = 9-22 males). (B) Silencing the CRNs does not increase mating drive after MTPA feeding (two-way ANOVA, n = 10-13 males). (C) Mutating GCE does not decrease courtship and prevents MTPA feeding from decreasing courtship (one-way ANOVA, n = 26-30 males). (D) RNAi knockdown of Met in the dopamine, NPF, and pCd neurons reduces mating drive (one-way ANOVA, n = 29-47 males). (E-F) RNAi knockdown of neither Met nor GCE in P1 (E) or CRN (F) neurons (bootstrap, E: n = 30-41 males, F: n = 20-29). (G-I) MTPA feeding does not increase CREB activity in the dopaminergic (G), P1 (H), or copulation-reporting (I) neurons (one-way ANOVA, G: n = 8 groups of 3 brains each, H: n = 7-8 groups, I: n = 8 groups each).

## REFERENCES

1. Abdou, M. a, He, Q., Wen, D., Zyaan, O., Wang, J., Xu, J., Baumann, A. a, Joseph, J., Wilson, T.G., Li, S., et al. (2011). Drosophila Met and Gce are partially redundant in transducing juvenile hormone action. Insect Biochem. Mol. Biol. 41, 938–945.

2. Argue, K.J., Yun, A.J., and Neckameyer, W.S. (2013). Early manipulation of juvenile hormone has sexually dimorphic effects on mature adult behavior in Drosophila melanogaster. Horm. Behav. 64, 589–597.

3. Baker, B.S., and Belote, J.M. (1983). Sex determination and dosage compensation in Drosophila melanogaster. Annu. Rev. Genet. 17, 345–393.

4. Bilen, J., Atallah, J., Azanchi, R., Levine, J.D., and Riddiford, L.M. (2013). Regulation of onset of female mating and sex pheromone production by juvenile hormone in Drosophila melanogaster. Proc. Natl. Acad. Sci. U. S. A. 110, 18321–18326.

5. Boccardo, F., Rubagotti, A., Barichello, M., Battaglia, M., Carmignani, G., Comeri, G., Conti, G., Cruciani, G., Rubagotti, A., Barichello, M., et al. (1999). Bicalutamide monotherapy versus flutamide plus goserelin in prostate cancer patients: results of an Italian Prostate Cancer Project study. J. Clin. Oncol. 17, 2027–2038.

6. Boutros, C.L., Miner, L.E., Mazor, O., and Zhang, S.X. (2017). Measuring and Altering Mating Drive in Male Drosophila melanogaster. J. Vis. Exp. e55291.

7. Bownes, M., and Rembold, H. (1987). The titre of juvenile hormone during the pupal and adult stages of the life cycle of Drosophila melanogaster. Eur. J. Biochem. 164, 709–712.

8. Burgess, C.R., Ramesh, R.N., Sugden, A.U., Levandowski, K.M., Minnig, M.A., Fenselau, H., Lowell, B.B., and Andermann, M.L. (2016). Hunger-Dependent Enhancement of Food Cue Responses in Mouse Postrhinal Cortex and Lateral Amygdala. Neuron 91, 1154–1169.

9. Chen, T.-W., Wardill, T.J., Sun, Y., Pulver, S.R., Renninger, S.L., Baohan, A., Schreiter, E.R., Kerr, R. a, Orger, M.B., Jayaraman, V., et al. (2013). Ultrasensitive fluorescent proteins for imaging neuronal activity. Nature 499, 295–300.

10. Clowney, E.J., Iguchi, S., Bussell, J.J., Scheer, E., and Ruta, V. (2015). Multimodal Chemosensory Circuits Controlling Male Courtship in Drosophila. Neuron 87, 1036–1049.

11. Coolen, L.M., and Wood, R.I. (1998). Testosterone stimulation of the medial preoptic area and medial amygdala in the control of male hamster sexual behavior: Redundancy without amplification. Behav. Brain Res. 98, 143–153.

12. Davidson, J.M. (1966). Characteristics of sex behaviour in male rats following castration. Anim. Behav. 14, 266–272.

13. Dietzl, G., Chen, D., Schnorrer, F., Su, K.-C., Barinova, Y., Fellner, M., Gasser, B., Kinsey, K., Oppel, S., Scheiblauer, S., et al. (2007). A genome-wide transgenic RNAi library for conditional gene inactivation in Drosophila. Nature 448, 151–156.

14. Durant, W. (1954). Our Oriental Heritage: The Story of Civilization.

15. Fan, P., Manoli, D.S., Ahmed, O.M., Chen, Y., Agarwal, N., Kwong, S., Cai, A.G., Neitz, J., Renslo, A., Baker, B.S., et al. (2013). Genetic and neural mechanisms that inhibit Drosophila from mating with other species. Cell 154, 89–102.

16. Feng, Y., Ueda, A., and Wu, C.-F. (2004). A Modified Minimal Hemolymph-Like Solution, HL3.1, for Physiological Recordings At the Neuromuscular Junctions of Normal and Mutant Drosophila Larvae. J. Neurogenet. 18, 377–402.

17. Freund, K. (1980). Therapeutic Sex Drive Reduction. Acta Psychiatr. Scand. 62, 5–38.

18. Godlewski, J., Wang, S., and Wilson, T.G. (2006). Interaction of bHLH-PAS proteins involved in juvenile hormone reception in Drosophila. Biochem. Biophys. Res. Commun. 342, 1305–1311.

19. Inoue, S., Yang, R., Tantry, A., Davis, C., Yang, T., Knoedler, J.R., Wei, Y., Adams, E.L., Thombare, S., Golf, S.R., et al. (2019). Periodic Remodeling in a Neural Circuit Governs Timing of Female Sexual Behavior. Cell 179, 1–16.

20. Kallman, B.R., Kim, H., and Scott, K. (2015). Excitation and inhibition onto central courtship neurons biases Drosophila mate choice. Elife 4, e11188.

21. Kimura, K., Hachiya, T., Koganezawa, M., Tazawa, T., and Yamamoto, D. (2008). Fruitless and Doublesex Coordinate to Generate Male-Specific Neurons that Can Initiate Courtship. Neuron 59, 759–769.

22. Kohatsu, S., and Yamamoto, D. (2015). Visually induced initiation of Drosophila innate courtship-like following pursuit is mediated by central excitatory state. Nat. Commun. 6, 6457.

23. Kohatsu, S., Koganezawa, M., and Yamamoto, D. (2011). Female contact activates male-specific interneurons that trigger stereotypic courtship behavior in Drosophila. Neuron 69, 498–508.

24. Krey, L.C., and Mcginnis, M.Y. (1990). Time-courses of the appearance/disappearance of nuclear androgen + receptor complexes in the brain and adenohypophysis following testosterone administration/withdrawal to castrated male rats: Relationships with gonadotropin secretion. J. Steroid Biochem. 35, 403–408.

25. Lawrence, P.A., and Johnston, P. (1986). The muscle pattern of a segment of Drosophila may be determined by neurons and not by contributing myoblasts. Cell 45, 505–513.

26. Levin, E.R. (2009). Plasma membrane estrogen receptors. Trends Endocrinol. Metab. 20, 477–482.

27. Lin, H.-H., Cao, D.-S., Sethi, S., Zeng, Z., Chin, J.S.R., Chakraborty, T.S., Shepherd, A.K., Nguyen, C.A., Yew, J.Y., Su, C.-Y., et al. (2016). Hormonal Modulation of Pheromone Detection Enhances Male Courtship Success. Neuron 1–14.

28. Liu, Q., Tabuchi, M., Liu, S., Kodama, L., Horiuchi, W., Daniels, J., Chiu, L., Baldoni, D., and Wu, M.N. (2017). Branch-specific plasticity of a bifunctional dopamine circuit encodes protein hunger. Science 539, 534–539.

29. Long, X., Colonell, J., Wong, A.M., Singer, R.H., and Lionnet, T. (2017). Quantitative mRNA imaging throughout the entire Drosophila brain. Nat. Methods 14, 703–706.

30. Mirth, C.K., Tang, H.Y., Makohon-Moore, S.C., Salhadar, S., Gokhale, R.H., Warner, R.D., Koyama, T., Riddiford, L.M., and Shingleton, A.W. (2014). Juvenile hormone regulates body size and perturbs insulin signaling in Drosophila. Proc. Natl. Acad. Sci. U. S. A. 111, 7018–7023.

31. Nielsen, J., Hedeholm, R.B., Heinemeier, J., Bushnell, P.G., Christiansen, J.S., Olsen, J., Ramsey, C.B., Brill, R.W., Simon, M., Steffensen, K.F., et al. (2016). Eye lens radiocarbon reveals centuries of longevity in the Greenland shark (Somniosus microcephalus). Science (80-.). 353, 702–704.

32. Pan, Y., Meissner, G.W., and Baker, B.S. (2012). Joint control of *Drosophila* male courtship behavior by motion cues and activation of male-specific P1 neurons. Proc. Natl. Acad. Sci. U. S. A. 109, 10065–10070.

33. Parker, L., Gross, S., Beullens, M., Bollen, M., Bennett, D., and Alphey, L. (2002). Functional interaction between nuclear inhibitor of protein phosphatase type 1 (NIPP1) and protein phosphatase type 1 (PP1) in Drosophila: consequences of over-expression of NIPP1 in flies and suppression by co-expression of PP1. Biochem. J. 368, 789–797.

34. von Philipsborn, A.C., Liu, T., Yu, J.Y., Masser, C., Bidaye, S.S., and Dickson, B.J. (2011). Neuronal control of Drosophila courtship song. Neuron 69, 509–522.

35. Phoenix, C.H., Goy, R.W., Gerall, A.A., and Young, W.C. (1959). Organizing action of prenatally administered testosterone propionate on the tissues mediating mating behavior in the female guinea pig. Endocrinology 65, 369–382.

36. Putnam, S.K., Du, J., Sato, S., and Hull, E.M. (2001). Testosterone Restoration of Copulatory Behavior Correlates with Medial Preoptic Dopamine Release in Castrated Male Rats. Horm. Behav. 224, 216–224.

37. Richard, D.S., Applebaum, S.W., Sliter, T.J., Baker, F.C., Schooley, D. a, Reuter, C.C., Henrich, V.C., and Gilbert, L.I. (1989). Juvenile hormone bisepoxide biosynthesis in vitro by the ring gland of Drosophila melanogaster: a putative juvenile hormone in the higher Diptera. Proc. Natl. Acad. Sci. U. S. A. 86, 1421–1425.

38. Riddiford, L.M. (2008). Juvenile hormone action: A 2007 perspective. J. Insect Physiol. 54, 895–901.

39. Riddiford, L.M., Truman, J.W., Mirth, C.K., and Shen, Y. (2010). A role for juvenile hormone in the prepupal development of Drosophila melanogaster. Development 1126, 1117–1126.

40. Robinson, G.E. (1992). Division of Labor in Insect Societies. Encycl. Insects 37, 637–655.

41. Schulz, K.M., Molenda-Figueira, H.A., and Sisk, C.L. (2009). Back to the future: The organizational-activational hypothesis adapted to puberty and adolescence. Horm. Behav. 55, 597–604.

42. Simerly, R.B. (1989). Hormonal control of the development and regulation of tyrosine hydroxylase expression within a sexually dimorphic population of dopaminergic cells in the hypothalamus. Mol. Brain Res. 6, 297–310.

43. Sisk, C.L., and Zehr, J.L. (2005). Pubertal hormones organize the adolescent brain and behavior. Front. Neuroendocrinol. 26, 163–174.

44. Södersten, P., Damassa, D.A., and Smith, E.R. (1977). Sexual behavior in developing male rats. Horm. Behav. 8, 320–341.

45. Spieth, H.T. (1974). Courtship behavior in Drosophila. Annu. Rev. Entomol. 19, 385–405.

46. Tanenhaus, A.K., Zhang, J., and Yin, J.C.P. (2012). In vivo circadian oscillation of dCREB2 and NF-κB activity in the Drosophila nervous system. PLoS One 7, e45130.

47. Thistle, R., Cameron, P., Ghorayshi, A., Dennison, L., and Scott, K. (2012). Contact chemoreceptors mediate male-male repulsion and male-female attraction during Drosophila courtship. Cell 149, 1140–1151.

48. Wang, L., Han, X., Mehren, J., Hiroi, M., Billeter, J.C., Miyamoto, T., Amrein, H., Levine, J.D., and Anderson, D.J. (2011). Hierarchical chemosensory regulation of male-male social interactions in Drosophila. Nat. Neurosci. 14, 757–762.

49. Wigglesworth, V.B. (1936). The function of the Corpus Allatum in the Growth and Reproduction of Rhodnius prolixus (Hemiptera). Q. J. Miscroscopical Sci. 79, 91–119.

50. Wigglesworth, V.B. (1965). The juvenile hormone. Nature 208, 522–524.

51. Wijesekera, T.P., Saurabh, S., and Dauwalder, B. (2016). Juvenile hormone is required in adult males for drosophila courtship. PLoS One 11, 1–11.

52. Wilson, T.G., DeMoor, S., and Lei, J. (2003). Juvenile hormone involvement in Drosophila melanogaster male reproduction as suggested by the Methoprene-tolerant27 mutant phenotype. Insect Biochem. Mol. Biol. 33, 1167–1175.

53. Wu, M. V, Manoli, D.S., Fraser, E.J., Coats, J.K., Tollkuhn, J., Honda, S.-I., Harada, N., and Shah, N.M. (2009). Estrogen masculinizes neural pathways and sex-specific behaviors. Cell 139, 61–72.

54. Wu, Z., Autry, A.E., Bergan, J.F., Watabe-Uchida, M., and Dulac, C.G. (2014). Galanin neurons in the medial preoptic area govern parental behaviour. Nature 509, 325–330.

55. Yamamoto, R., Bai, H., Dolezal, A.G., Amdam, G., and Tatar, M. (2013). Juvenile hormone regulation of Drosophila aging. BMC Biol. 11, 85.

56. Yan, H., Opachaloemphan, C., Mancini, G., Rgen Liebig, J., Reinberg, D., Desplan, C., Yang, H., Gallitto, M., Mlejnek, J., Leibholz, A., et al. (2017). An Engineered orco Mutation Produces Aberrant Social Behavior and Defective Neural Development in Ants. Cell 170, 736–747.

57. Zhang, S.X., Rogulja, D., and Crickmore, M.A. (2016). Dopaminergic Circuitry Underlying Mating Drive. Neuron 91, 168–181.

58. Zhang, S.X., Miner, L.E., Boutros, C.L., Rogulja, D., and Crickmore, M.A. (2018). Motivation, perception, and chance converge to make a binary decision. Neuron 99, 1–13.

59. Zhang, S.X., Rogulja, D., and Crickmore, M.A. (2019). Recurrent Circuitry Sustains Drosophila Courtship Drive While Priming Itself for Satiety. Curr. Biol. 29, 3216–3228.e9.

